# Mutational and evolutionary dynamics of non-structural and spike proteins from variants of concern (VOC) of SARS-CoV-2 in India

**DOI:** 10.1101/2024.07.14.603481

**Authors:** Ankur Chaudhuri, Subhrangshu Das, Saikat Chakrabarti

## Abstract

Monitoring the genetic diversity and emerging mutations of SARS-CoV-2 still remains to be crucial in India. This study extensively analyzes the lineage dynamics, mutation screening, structural analysis, and phylodynamics of SARS-CoV-2 variants of concern (VOC) in India from October 2020 to September 2023. Predominant variants identified include alpha, beta, delta, and omicron, with delta and omicron making up 76.05% of sequenced genomes. The B.1.617.2 lineage of delta was the major contributor to COVID-19 cases before the rapid rise of omicron. Mutation screening of non-structural proteins (NSPs) and spike revealed distinct profiles for each VOC. Co-mutation patterns/networks of the most frequently observed mutations specific for each VOC were also identified and subsequently structural and energetic alteration of the co-mutants were analyzed using rigorous molecular dynamics simulations. Furthermore, comparative analysis of phylogenetic trees based on genomic and mutational data revealed that nsp1, nsp3, nsp4, nsp13, and nsp14 exhibit strongest association with the increased mutation load across the genome of SARS-CoV-2 in Indian population. Our comparative phylogenetic study also revealed that mutability patterns of nsp14 and spike have highest similarity supporting critical role of nsp14 for SARS-CoV-2 infectivity and persistence. This research provides a comprehensive overview of SARS-CoV-2 evolution in India.

## Introduction

The COVID-19 pandemic caused by SARS-CoV-2has caused more than 770 million cases of COVID-19 and more than 6.9 million deaths as of September 2023 [1]. After recording its first case in India on January 27, 2020, the country now has a total of about 45 million confirmed coronavirus cases with 0.55 million deaths as of September 2023 according to the COVID-19 Data Repository of the World Health Organization [2]. The genome of SARS-CoV-2 consists of 10 open reading frames (ORFs), which encode a total of 27 proteins [3]. Among these, ORF1ab encodes 16 non-structural proteins (NSPs) [4–6], while the structural proteins comprise spike (S), envelope (E), membrane (M), and nucleocapsid (N) proteins [6–7]. To enhance research efforts and global surveillance concerning SARS-CoV-2 variants, the World Health Organization (WHO) has categorized genetic lineages of the virus into three distinct classifications: variants of concern (VOC), variants of interest (VOI), and variants under monitoring (VUMs) [8]. At present, five variants and their descendant lineages have been designated as VOC. Among them, the Alpha (B.1.1.7) [9], Beta (B.1.351) [10], Gamma (P.1) [11], and Delta (B.1.617.2) [12] variants were previously predominant, while the Omicron (B.1.1.529) variant and its sub-lineages [13–14] are currently prevalent. Presently, no VOC are actively circulating, though certain Omicron sub-variants, such as XBB.1.5, XBB.1.16, EG.5, BA.2.86, and JN.1 are under monitoring [15–18].

The structural and non-structural proteins (NSPs) of SARS-CoV-2 play important roles in its infectivity and replication machinery. For instance, they perform a variety of functions, such as nsp3 and nsp5, which are viral proteases involved in processing the polyproteins [19–21]. Nsp12, working alongside Nsp7 and Nsp8 as co-factors, operates RNA synthesis process [22–23]. Nsp13 acts as a helicase and nsp14 as an exoribonuclease activity for proofreading [24–25]. Other NSPs are involved in host cell interaction and immune suppression [26–28]. The structural proteins play vital roles in viral interaction with the host cell’s receptor, viral entry, morphogenesis, assembly, membrane fusion, and release of virion particles [6–7]. These proteins are crucial for the virus’s ability to replicate efficiently and avoid host defences, making them targets for antiviral drug development.

Since the onset of the SARS-CoV-2 pandemic, numerous studies have highlighted the role of mutations at both the genomic and protein levels in driving evolutionary changes [29–30]. These mutations are crucial for understanding the virus’s ability to successfully invade and infect host cells [31–32]. To address this, we conducted an examination of 2,85,321 SARS-CoV-2 genomes of Indian origin, with 77,516 sequences selected for in-depth scrutiny. To the best of our knowledge, such detailed comparative investigation regrouping all variants of concern (VOC) utilizing lineage dynamics, time-dependent mutation screening, co-mutation analyses, genome-based and mutation based phylodynamics and structure-based simulation is still lacking to understand the functional impact of these mutations on the SARS-CoV-2 VOC specific mechanism of action.

Here, we investigated the mutation frequency (MF) of NSPs and spike in VOC of SARS-CoV-2 and explored their lineage dynamics across different time scales. We employed a stepwise approach of phylogenetic analyses in combination with genome-based and mutation-based approach by utilizing prevalent lineages associated with each VOC. Additionally, we analyzed the co-mutation patterns by combining sets of 3 to 10 mutations within each VOC. By analyzing the frequency and distribution of co-mutated sites, we aimed to elucidate potential association among various mutations and their collective impact on viral evolution and adaptation. This approach enabled us to uncover networks of co-evolving mutations within VOC. Furthermore, we present results from structure-based in silico modeling and a full-atomistic molecular dynamics (MD) simulation of selected co-mutational cliques from each VOC. Taken together, we facilitated a comprehensive understanding of the molecular determinants underlying the enhanced transmissibility and pathogenicity observed with the variant of concern of SARS-COV-2 prevalent in India.

## Materials and Methods

### Lineage dynamics

All the metadata of variants of concern (VOC) of SARS-CoV-2 were collected from the EpiCoV database of the Global Initiative on Sharing All Influenza Data (GISAID) platform [33]. The metadata includes information on the genomic sequence of the variant, as well as epidemiological data related to the sample, such as the location and date of sample collection. The dataset was retrieved by filtering the time period from 1^st^ October 2020 to 30^th^ September 2023 using the keywords “hCoV-19”, “India” and “human”. Sequences with genomes > 29,000 bp and <1% Ns (undefined bases) were considered as complete and high-coverage sequences respectively. Only complete and high-coverage sequences from India were selected. Gamma VOC was no included due to very less numbers of genomes was submitted till 30^th^ September 2023 possessing complete metadata. Figure S1 provides a flowchart of the overall methodology and the following sections briefly describe the methods.

### Mutation screening and frequency calculation

Using the metadata of individual VOC of SARS-CoV-2, the mutation frequency of each amino acid of spike and non-structural proteins (NSP1-16) was calculated and analyzed. To track the mutational dynamics of individual amino acids of each protein of different VOC, the timeline was divided into several periods like Oct’20 - Dec’20, Jan’21 - Mar’21, Apr’21 - Jun’21, Jul’21 - Sep’21, Oct’21 - Dec’21, Jan’22 - Mar’22, Apr’22 - Jun’22, Jul’22 - Sep’22, and Oct’22 - Sep’23, respectively. Frequencies of amino acid mutations at specific positions were computed directly from the metadata using in-house Python scripts. Mutation frequency was calculated by dividing the number of occurrences of each amino acid mutation by the total number of sequences.

### Phylodynamics and comparative analysis

Phylodynamics is a field that combines principles from phylogenetics and epidemiology to study the dynamics of pathogen evolution within host populations. In this study, we examined the leading ten lineages of SARS-CoV-2, including B.1.1.7 representing alpha [9], B.1.351 representing beta [10], B.1.617.2, AY.112, and AY.127 representing delta, as well as BA.1.1, BA.2, BA.2.10, BA.2.38, and XBB.1.16 representing omicron [34] VOC. The selection of lineages was based on their prevalence, diversity, and significance in the spread of the virus within the Indian population. Altogether, 500 genome sequences representing ten different lineages of variants of concern (VOC) were obtained from the GISAID database, with 50 sequences randomly retrieved from each specific lineage. The combined dataset of all the genomes were aligned using the MUSCLE algorithm [35] implemented in the MEGA X tool [36]. The phylogenetic tree was constructed using the UPGMA method [37] in MEGA X with 1,000 bootstraps. Alternatively, a different phylogenetic tree was built by utilizing mutational data from all ten distinct lineages of the VOC. The web tool Phylo.io [38] and Visual Tree cmp [39] were utilized to perform the pairwise comparisons between genome-based and mutation-based phylogenetic trees, evaluating the similarity or divergence in their tree topology. The phylogenetic trees were visualized using the Interactive Tree Of Life (iTOL; version 6.1.1) online tool [40].

### Co-mutation analysis

Co-mutation analysis provides valuable perspectives into the evolutionary dynamics and molecular mechanisms driving SARS-CoV-2 infection via exploring the simultaneous occurrence of mutations across various VOC within the viral genome. To identify the most prevalent co-mutation patterns among individual VOC, we categorized prevalent mutations into eight groups or cliques considering 3-10 co-mutations in spike and NSPs for each viral genomes from respective VOC. The dominant co-mutation from each category within individual VOC was visualized in a network representation generated using the Cytoscape software [41]. The five-member clique containing five co-mutations was most prevalent in every VOC. Therefore, we selected a five-member clique from each VOC for further structural analysis.

### Molecular modeling and dynamics simulation

Initially, wild-type structures of NSPs and spike proteins were retrieved from the PDB database [42]. Each mutated structure of an individual NSP and spike was generated by MODELLER 9v10 [43]. From the resultant 100 models, the best model was selected on the basis of the DOPE score [44]. In this study, some of the NSPs and spike mutant proteins were also generated using the neural network based algorithm AlphaFold 2 colab [45]. AlphaFold computes the pLDDT score [46] and pTM score [47] to indicate the accuracy of a prediction. The top prediction as ranked by pLDDT was used in further analysis.

All-atom molecular dynamics (MD) simulations were used to capture the structural and energetic properties of the spike and NSP mutants. We selected a five-membered clique from each VOC for molecular dynamics simulation study, aiming to analyze the structural alteration of resulting from the multiple co-mutations within a SARS-CoV-2 variant. GROMACS v 2018 [48] and AMBER 99SB force field [49] were used for theMD simulations. Protein system was solvated in a cubic box with a TIP3P water model [50] by maintaining periodic boundary conditions (PBC) [51] through the simulation process. Sodium and chloride ions were added to neutralize all the systems. Each system was energy minimized using the steepest descent algorithm [52] until the maximum force was smaller than 1000 kJ/mol/nm. This was done to remove any steric clashes in the system. Each system was equilibrated with a 200 ps isothermal-isochoric ensemble, NVT followed by a 200 ps isothermal-isobaric ensemble NPT. The two types of ensemble of equilibration methods stabilized systems at 310 K and 1 bar pressure. The Berendsen thermostat [53] and Parrinello-Rahman [54] methods were applied for temperature and pressure coupling, respectively. Particle Mesh Ewald (PME) method [55] was used for the calculations of the long-range electrostatic interactions and the cut-off radii for Van der Waals and coulombic short-range interactions were set to 0.9 nm. The Linear Constraint Solver (LINCS) constraints algorithm [56] was used to fix the lengths of the peptide bonds and angles. All the systems were subjected to MD simulations for 100 ns each. The resulting MD trajectories were utilized through the inbuilt tools of GROMACS for analysis purposes. The subsequent analyses were performed using VMD [57], USCF Chimera [58], Pymol [59], and xmgrace [60], respectively. DSSP analysis [61] was performedto inspect secondary structural element formations during MD simulations.

### Clustering of conformations for ensemble generation and Essential Dynamics

To evaluate the dominant conformation acquired by proteins from each VOC throughout the MD simulations, clustering analysis was conducted. The entire molecular dynamics trajectories underwent RMSD-based clustering using the ‘gmx cluster’ tool to explore the conformational diversity among the ensemble of protein structures. To further investigate the dynamic behavior of the proteins associated with each VOC, Essential Dynamics (ED) was performed [62]. ED is a technique that identifies the essential motions present in the MD trajectory of the targeted protein molecules, which are typically correlated and biologically significant for their functions. The set of eigenvectors and eigenvalues was computed by diagonalizing the covariance matrix. The covariance matrix and eigenvectors were analyzed using the ‘gmx cover’ and ‘gmxanaeig’ tools, respectively.

## Results and Discussions

### Prevalence of SARS-CoV-2 VOC in India from 2020-2023

From 1^st^ October 2020 to 30^th^ September 2023, a total of 285,321 cases involving SARS-CoV-2 variants were detected in India. Following the application of filters, our study focused on 77,516 cases. Of those samples, alpha was confirmed in 2.70% (n= 2094/77516), beta in 0.18% (n= 146/77516), delta in 54.13% (n= 41961/77516), and omicron in 21.92% (n= 16992/77516). The remaining 21.87% (n= 16353/77516) deposited genomes were from other/mixed variant strains. The prevalence of each VOC across every time-period is shown in Figure 1A. Different variants rose to dominance at different times and during different infection waves across the country. The first VOC to emerge was the alpha variant, first identified in Oct ‘20. In the Oct’20 - Mar’21 study period, the alpha variant significantly increased, growing from 1.96% (n = 60/3047) to 15.28% (n = 1064/6981). It became the dominant strain in the country till Mar ‘21. Compared to the original Wuhan strain of the virus, the alpha variant was more contagious and more likely to result in severe disease. Within this time scale, beta variants also emerged (prevalence: 1.16%; n= 81/6981). From Apr ‘21 - Jun’21, alpha and beta variants were reduced to 5.72% (n= 960/16779) and 0.38% (n = 65/16779), respectively. Between Oct’20 and Dec’20, the delta variant was identified in parallel with the alpha variant. This variant rapidly replaced previous VOC leading to a drastic surge in Covid infection cases around the world. The most significant increase in its prevalence in India was observed during Jul ‘21 - Sep’21 amounting 94.02% (n= 12695/13502) of the deposited genomes and remained dominant until Dec’21. Subsequently, it was reduced to 4.22% (n= 416/9855) in Mar ‘22. The delta variant was the most dominant strain in India during the third wave of COVID-19. The omicron variant was first identified in Dec’21. This variant started new global waves of infection. Sequences clustering to omicron had been detected as early as Dec’21 (8.51%, n = 1587/18638) but became dominant during Sep’22 (47.77%, n= 1243/2602). The most significant increase in its prevalence was observed in Oct ‘22 - Sep’23 reaching to 99.22% (n= 2756/2772). The duration of a VOC prevalence can change based on several variables, such as the variant’s transmissibility, the success of public health initiatives, and the vaccination rates in various nations. However, the COVID-19 pandemic has been significantly impacted by all of the VOC that have been discovered in India, increasing cases, hospital admissions, and fatalities. Overall, the delta (54.13%) and omicron (21.92%) constitute the top viral variants within the sequenced SARS-CoV-2 genomes in India over the whole period.

**Figure 1:**
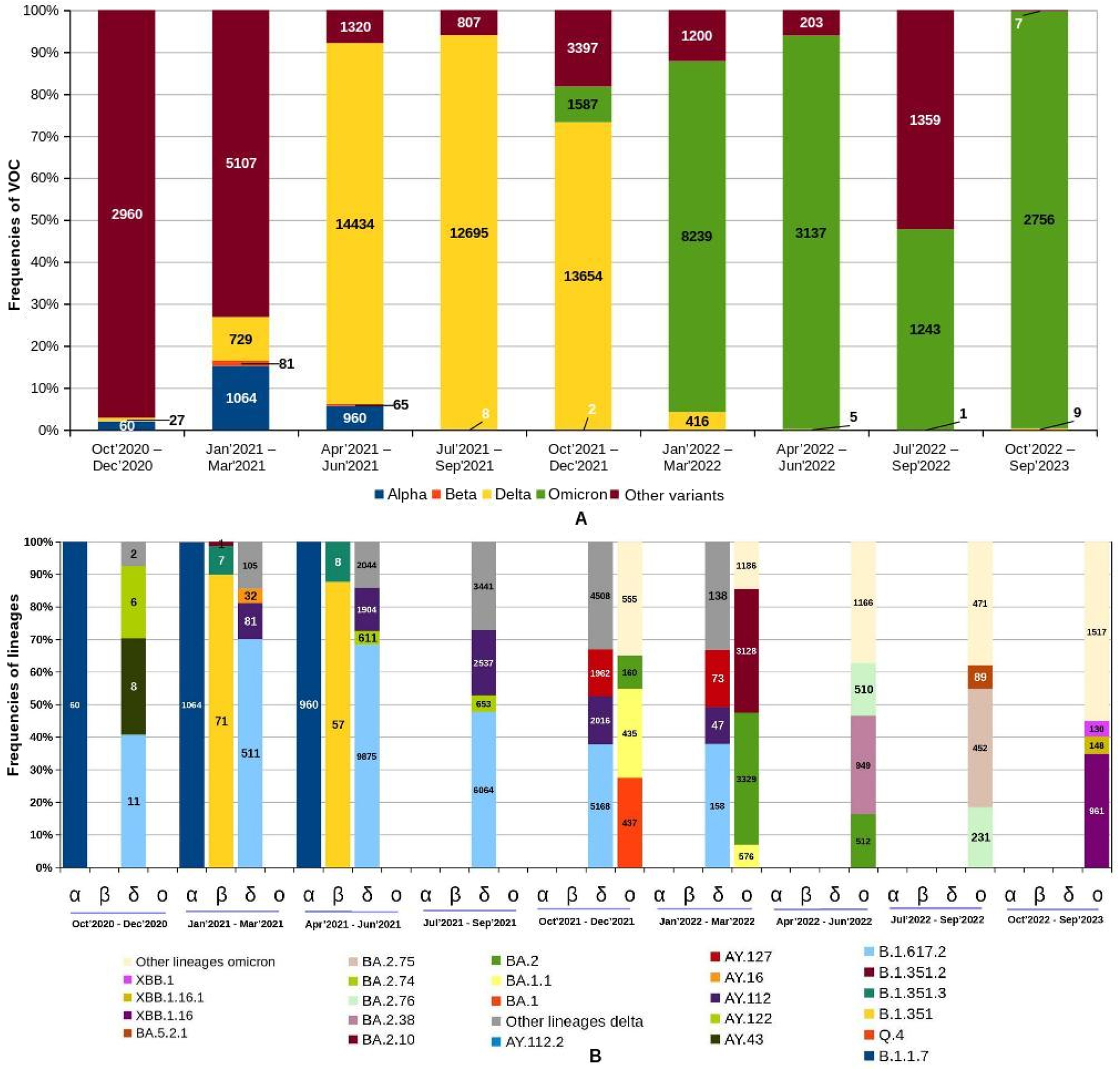
Dynamics and prevalence of VOC: A. Trends in the prevalence of major variants circulating in India from October 2020 to September 2023. The *Y*-axis shows the percentage distribution of VOC across the various time periods (*X*-axis) while different colors represent the VOC. B. Representations of the top three/four lineages within the VOC in different time periods spanning from October 2020 to September 2023.

### Lineage dynamics of VOC of SARS-CoV-2

Pangolin nomenclature tool was used to assign all sequences to corresponding lineages in order to investigate the lineage dynamics across the different outbreak waves. We identified 432 lineages (pangolin nomenclature) of VOC of SARS-CoV-2 for the entire duration of the study period in India. Out of these lineages, a total of 2, 3, 148, and 279 different lineages were identified for alpha (0.46%, n= 2/432), beta (0.69%, n=3/432), delta (34.25%, n= 148/432) and omicron (64.58%, n=279/432), respectively. Following analyses focused on the different lineages that contributed to the surges during the several waves of the SARS-CoV-2 infection in the Indian population (Figure 1B). During Oct’20 - Jun’21, the dominant lineage of the alpha variant was B.1.1.7. A steep reduction in the B.1.1.7 lineage was observed in Jun ‘21. In the study period of Jan’21 - Jun’21, three different lineages of the beta variant were circulating in the Indian population. These lineages were B.1.351, B.1.351.2, and B.1.351.3. The most predominant lineage was B.1.351 (88.88 %, n= 128/144).

VOC delta was first detected in Dec’20, concordant with the emergence in India, and rapidly dominated transmission. We detected 148 lineages (pangolin nomenclature) in 41,961 genomes of delta variant sampled for the entire duration of the study period. Out of these lineages, the top four prevalent lineages were B.1.617.2 (51.92%, n= 21789/41961), AY.112 (15.69%, n= 6586/41961), AY.127 (6.52%, n= 2739/41961), and AY.122 (4.27%, n= 1792/41961). The time period of Jan’21 - Jun’21 serves as a transitional period between three major variants (alpha, beta, and delta). Alternatively, the transmission of alpha and beta declined when delta was introduced and emerged. The dynamics of the delta variant change over every three months. During the first appearance of the delta variant (Oct’20 - Dec’20), the top three lineages such as B.1.617.2 (40.74%, n= 11/27), AY.43 (29.62%, n= 8/27), and AY.122 (22.22%, n= 6/27) were detected. Between Jan’21 - Mar’21, the most prevalent circulating lineages were B.1.617.2 (70.09%, n= 511/729), AY.112 (11.11%, n= 81/729), and AY.16 (4.38%, n= 32/729). During Apr’21 - Jun’21, B.1.617.2 (68.41%, n= 9875/14434) was the predominant followed by AY.112 (13.19%, n=1904/14434) and AY.122 (4.23%, n= 611/14434). These three predominant lineages continued to be detected in the study period of Jul’21 - Sep’21. Between Oct’21 - Dec’21, B.1.617.2 (37.84%, n= 5168/13654), AY.112 (14.76%, n= 2016/13654), and AY.127 (14.36%, n= 1962/13654) were the top three prominent lineages circulating in the Indian population. Of all lineages of the delta variant, B.1.617.2 lineage was common in all time-period. It is characterized broadly as the most dominant emerging lineage, replacing most lineages, and causing most COVID-19 cases in India from its emergence Dec ‘20. In a short period, the delta variant of SARS-CoV-2 emerged as a major strain across Indian states and was transmitted to more than 40 countries, crippling health systems globally. A sharp decrease in the different lineages of the delta variant was observed in the time period of Jan’22 - Mar’22.

Omicron was first identified in India in December 2021; it quickly became dominant in India. It was declared a VOC by the WHO on November 26, 2021. The time period of Dec’21 - Feb’21 serves as a transitional period between the delta and omicron variants. Due to its substantial growth advantage over the delta, the omicron variant rapidly replaced the delta variant globally. Omicron induced less severe disease than the delta variant and resulted in lower hospitalization and mortality rates. We identified 279 lineages (pangolin nomenclature) in 16,992 genomes of omicron variant deposited the entire duration of the study period. Out of these lineages, the top four prevalent lineages were BA.2 (23.96%, n= 4072/16992), BA.2.10 (19.62%, n= 3335/16992), BA.2.38 (6.16%, n= 1048/16992), and BA.1.1 (5.94%, n= 1011/16992). In Dec’21, the top three prevalent circulating lineages were BA.1 (27.53%, n= 437/1587), BA.1.1 (27.41%, n= 435/1587), and BA.2 (10.08%, n= 160/1587). Between Jan’22 - Mar’22, BA.2 (40.50%, n= 3329/8219) and BA.2.10 (38.05%, n= 3128/8219) were the most circulating lineages followed by BA.1.1 (7.00%, n= 576/8219). During the study period of Apr’22 - Jun’22, different sub-lineages of the BA.2 evolved. The most dominant was BA.2.38 (30.25%, n= 949/3137) followed by BA.2.74 (16.32%, n= 512/3137) and BA.2.76 (16.25%, n= 510/3137). As of Jul’22 - Sep’22, BA.2.75 (35.39%, n= 452/1277), BA.2.76 (18.08%, n= 231/1277), and BA.5.2.1 (6.96%, n= 89/1277) sub-lineages accounted for a cumulative prevalence of approximately 60.43%. During the study period of Oct’22 - Sep’23, the most prevalent lineages were XBB.1.16 (34.86%, n= 961/2756), XBB.1.16.1 (5.37%, n= 148/2756), and XBB.1 (4.71%, n= 130/2756).

### Mutation screening of non-structural proteins (NSPs 1-16)

The mutational dynamics of non-structural proteins in VOC of SARS-CoV-2 have significant implications for disease severity and viral adaptation. To investigate the implications of mutations on the transmission, infection, and virulence of SARS-CoV-2, we focus on the high-frequency or top mutations that represent the most common characteristics of the VOC of SARS-CoV-2 in India. For each NSP, mutations observed with frequency of 5% or more were taken into consideration. Higher (≥ 5%) frequencies of mutations in NSPs from the Indian samples were observed especially for nsp1, nsp2, nsp3, nsp4, nsp5, nsp6, nsp9, nsp12, nsp13, nsp14, and nsp15, respectively. The analysis of the VOC of the SARS-CoV-2 revealed that no mutation was observed with more than 90% frequency in nsp7, nsp8, nsp9, nsp10, nsp11 and nsp16 genes/proteins. We calculated and analyzed the mutation frequencies based on three categories: 1) mutation frequency of all variants (alpha, beta, delta, and omicron) in total time-scale (Oct ‘20 - Sep’23), 2) mutation frequency of top four lineages of delta and omicron variants in total time-scale, 3) mutation frequency of top lineage of delta and omicron variants according to individual time-scale. B.1.617.2 was the top lineage of the delta variant at each specific time scale during the duration of the whole study period. Throughout the entire study period, the top four lineages of the delta VOC were B.1.617.2, AY.112, AY.127, and AY.122. The mutation dynamics of the omicron VOC are distinctive. The top lineages from Oct’21 - Dec’21, Jan’22 - Mar’22, Apr’22 - Jun’22, and July’22 - Sep’22 were BA.1, BA.2, BA.2.38, and BA.2.75, respectively. The top four lineages of the omicron variation for the whole study period were BA.2, BA.2.10, BA.2.38, and BA.1.1. Following section briefly describe the mutational pattern within each NSPs and spike.

#### Nsp1 protein

Nsp1 is associated with the regulation of host translation and the viral infection cycle [26]. In the Indian population, the nsp1 gene predominantly displayed the S135R and K47R mutations, which are most commonly linked to the omicron variant (Figure 2A). The prevalence of these mutations in the omicron variant was 85.53% for S135R and 13.89% for K47R (Figure 2B). While the mutation frequency of S135R was below 1% in BA.1.1, it exceeded 90% in BA.2, BA.2.10, and BA.2.38 (Figure 2C). Interestingly, S135R mutation frequency surpassed 90% in all major lineages over time, except in BA.1. On the other hand, K47R was predominantly present at 97.81% of the XBB.1.16 lineage (Oct’22 - Sep’23) of the omicron variant (Figure 2D). This distinct mutation at positions 47 and 135 has a significant impact on the overall interaction and function of the N and C-terminal domains of the nsp1 protein. Notably, S135R and K47R are unique mutations in the omicron variant, absent in the alpha, beta, and delta variants.

**Figure 2:**
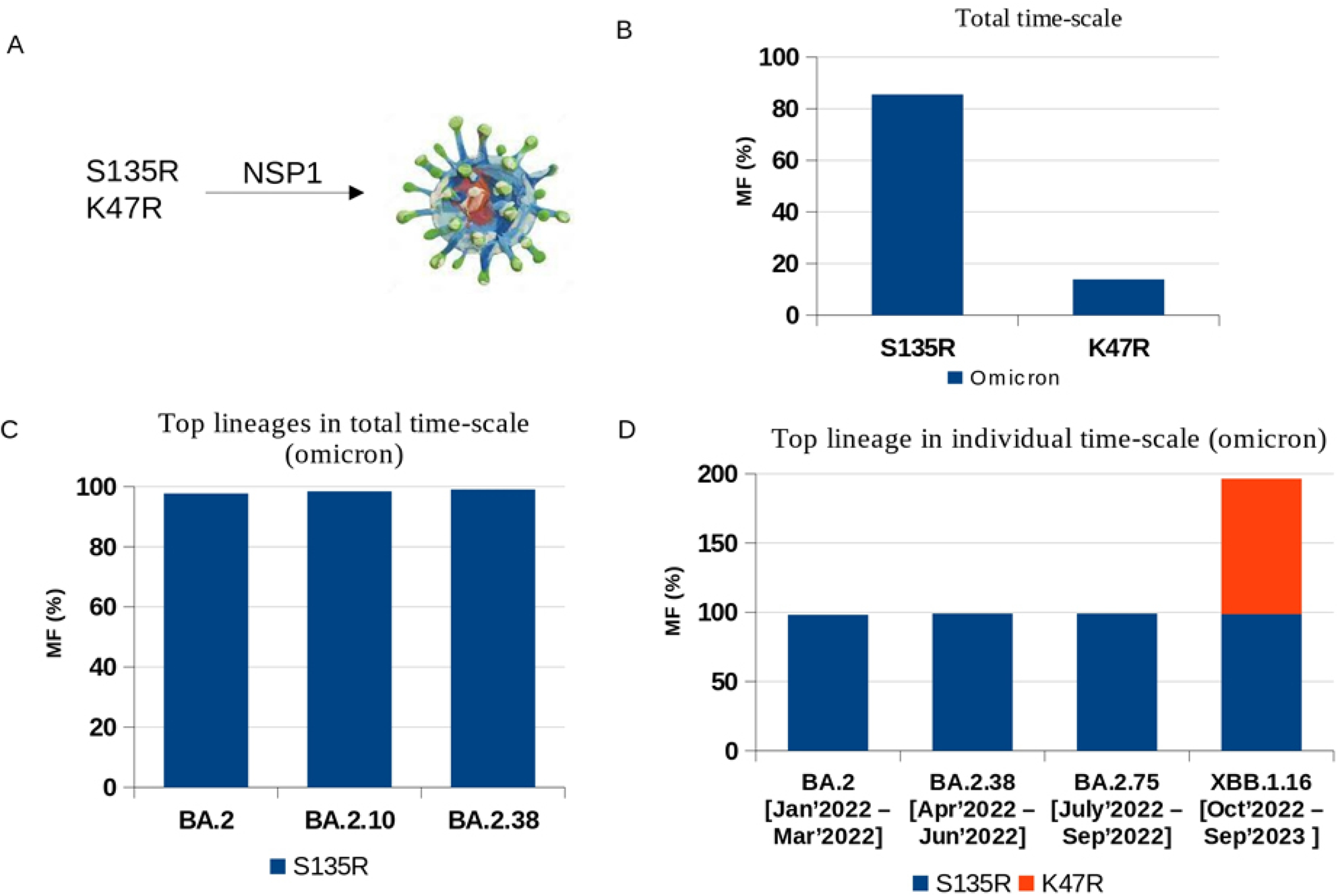
Mutational dynamics of nsp1: A. Mutated amino acid; B. Mutation frequency in total time-scale; C. Mutation frequency of top four lineages of omicron variant; D. Mutation frequency of top lineage in individual time periods.

#### Nsp2 protein

Nsp2 plays a crucial role in diverse biological processes such as host immune regulation, viral replication, endosomal transport, and mitochondrial biogenesis [27]. Mutational hotspots identified in nsp2 include T85I, P129L, L550F, and L580I (Figure S2A). Among these, the alpha variant exhibited the highest frequencies for L550F (11.6%) and T550I (18.16%). The beta variant predominantly featured the T85I mutation at a frequency of 93.29%. In the delta variant, P129L emerged as the most prevalent nsp2 mutation, with a frequency of 31.45% (Figure S2B). Remarkably, the B.1.617.2 lineage within the delta VOC uniquely retained the P129L mutation (Figure S2C), and this mutation persisted over the observed time period (Figure S2D). Notably, the omicron variant in the Indian population displayed nsp2 mutation rate of less than 1%.

#### Nsp3 protein

The papain-like protease (PLpro) domain is responsible for cleaving nsp3 from polyproteins pp1a and pp1ab [19]. Due to its large size, nsp3 exhibits distinct mutations in various VOC (Figure 3A). The top three mutations in alpha, each occurring with a frequency of above 30%, were T183I, A890D, and I1412T. Beta displays S794L and K837N mutations with frequencies of 22.56% and 37.15%, respectively. Three mutations with frequency greater than 60% were present in the delta variant: P1228L, A488S, and P1469S. The P822L mutation is associated with beta and delta, with frequencies of 9.14% and 32.48%, respectively. In the omicron variant, T24I, K38R, G489S, and A1892T have frequencies of 80.67%, 17.26%, 81.04%, and 17.24%, respectively (Figure 3B).

**Figure 3:**
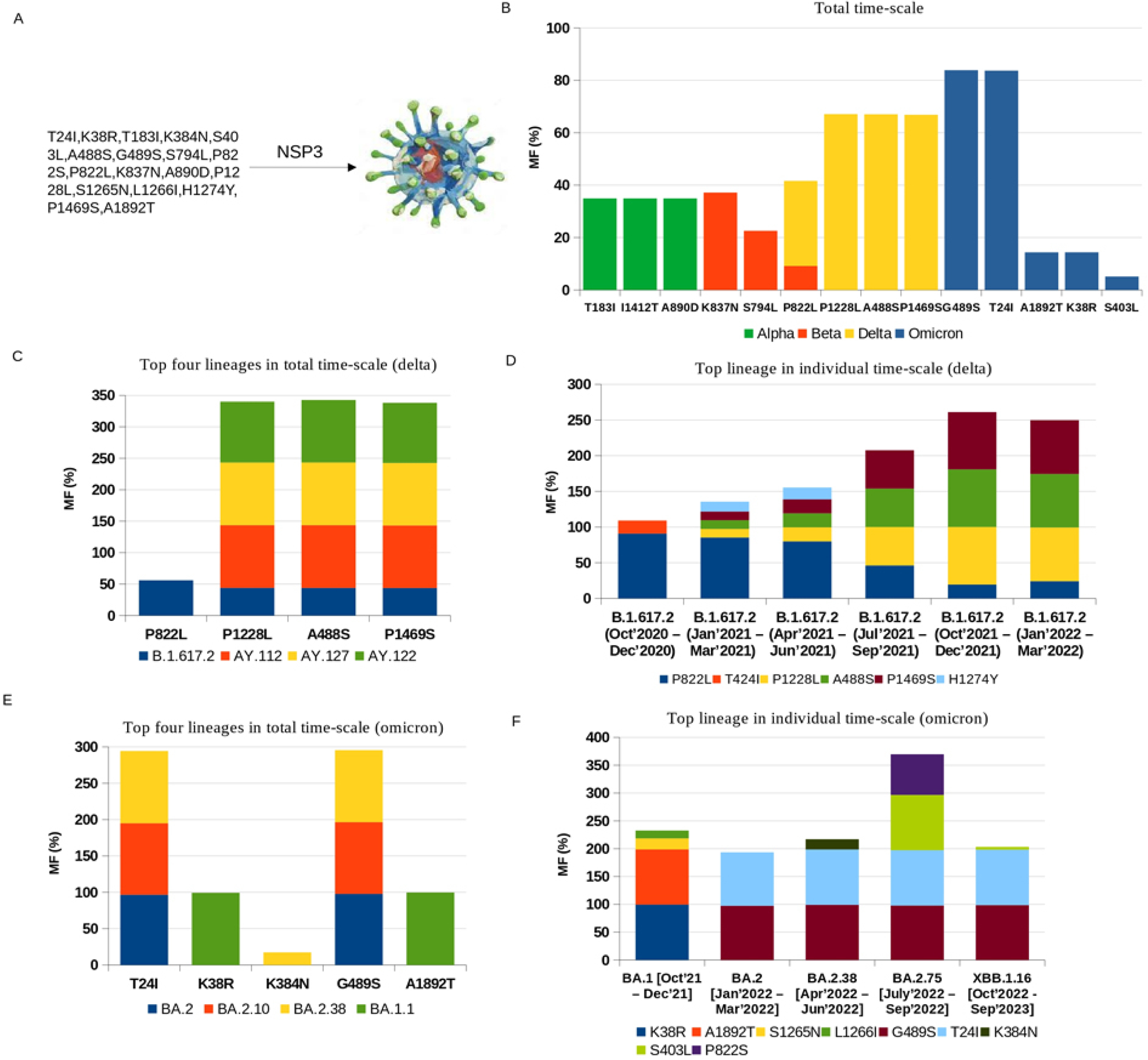
Mutational dynamics of nsp3: A. Mutated amino acids; B. Mutation frequency in total time-scale; C. Mutation frequency of top four lineages of delta variant; D. Mutation frequency of top lineage in individual time-scale of delta variant; E. Mutation frequency of top four lineages of omicron variant; F. Mutation frequency of top lineage in individual time-scale of omicron variant.

Throughout the study, the top four lineages of the delta VOC were B.1.617.2, AY.112, AY.127, and AY.122. The P822L mutation was specific to B.1.617.2, with a frequency of 55.95%. A488S, P1228L, and P1469S were present in B.1.617.2 at 43%, while exceeding 95% in the other three lineages. B.1.617.2 consistently dominated the delta VOC throughout the entire study period (Figure 3C). In B.1.617.2, the P822L mutation decreased from 90.09% to 24.05%, while A488S and P1228L increased from 12.32% to 75.05% and 11.93% to 75.31%, respectively (Figure 3D).

The top four omicron lineages were BA.2, BA.2.10, BA.2.38, and BA.1.1, all showing >98% detection of T24I and G489S mutations. K38R and A1892T were specific to BA.1.1, with frequencies >99% (Figure 3E). The dynamics of the top omicron lineage changed over time, with unique mutations such as K38R, A1892T, S1265N, and L1266I detected only in the BA.1 sub-lineage. In the BA.1 sub-lineage, K38R and A1892T had frequencies of 99.54% and 99.08%, while T24I and G489S exceeded 90% in BA.2, BA.2.38, BA.2.75, and XBB.1.16. The P822L mutation in the B.1.617.2 sub-lineage was replaced by P822S with a 73% frequency in the BA.2.75 sub-lineage of the omicron VOC. Both BA.2.75 and BA.2.38 exhibited S403L and K384N, with frequencies of 99.33% and 18.44%, respectively (Figure 3F).

#### Nsp4 protein

Nsp4, a protein spanning the membrane, plays a crucial role for the assembly of the viral replication-transcription complex (RTC) [28]. Nsp4 mutations were found to be less than 1% in the alpha VOC among the Indian population. Notably, in the beta variant, K8N and I379V exhibited specific mutation frequencies of 17.0% each. In the delta variant, A446V and V167L were observed with frequencies of 29.5% and 51%, respectively. The T492I mutation was shared by both delta and omicron, with frequencies of 66.85% and 98.85%, respectively. In the omicron variant, T327I, L438F, and L264F had frequencies of 78.54%, 77.52%, and 81.45%, respectively (Figure S3A-S3B). Throughout the study period, the predominant lineage of the delta VOC was B.1.617.2. In this sub-lineage, the A446V mutation frequency decreased from 72.72% to 19.62%, while T492I and V167L mutation frequencies increased from 12.32% to 80.39% and 13.01% to 69.67%, respectively (Figure S3D). These two mutations were present in all top four sub-lineages of the delta variant. The A446V mutation was specific to B.1.617.2, with a frequency of 50.22% (Figure S3C).In the omicron variant, T327I, L264F, and L438F were the most prevalent mutations, with frequencies exceeding 98% in all top sub-lineages, except BA.1.1 (Figure S3E). The T492I mutation was consistently present in all top four lineages and the top lineage across the study period, with a frequency exceeding 98% (Figure S3F). These findings suggest a potentially significant role of the T492I mutation in the transmission and fitness of SARS-CoV-2.

#### Nsp5 protein

Nsp5 plays a crucial role in the processing of viral polypeptides and serves as the primary protease (Mpro) in SARS-CoV-2 [20]. AnalyzingMpro metadata revealed prevalent mutations, namely K90R, P132H, and P184 - T196del (Figure S4A). The Mpro mutation frequency in the alpha VOC within the Indian population was less than 0.5%. K90R was exclusive to beta VOC, with a frequency exceeding 90%. The delta VOC (B.1.617.2) dominant in the latter part of 2021, did not exhibit prevalent (≥ 5%) missense mutations in the Mpro polyprotein region. The P132H mutation, prevalent over 90%, was notable in the omicron variant (Figure S4B). This mutation was consistently present in the top four lineages and the primary lineage over time, affecting domain II and enzyme activity (Figure S4C and S4D). The P184 - T196 deletion mutation was specific to the BA.2 sub-lineage of the omicron VOC, with a 15.65% frequency (Figure S4D). Elimination of this loop region in the sub-lineage contributed to the stability of the omicron variant.

#### Nsp6 protein

Nsp6, a crucial component of the viral genome replication and transcription machinery, collaborates with nsp3 and nsp4 to redirect and reorganize host cell membranes, leading to the formation of double-membrane vesicles (DMVs) [22]. A sequential deletion of three amino acids (S106-, G107-, and F108-) was consistently observed in all variants of concern (VOC) except the delta variant. In the alpha, beta, and omicron variants, the frequency of the ΔSGF mutation was 48%, 29%, and 14%, respectively (Figure S5A-S5B). Notably, the F108del mutation was replaced by F108L, with frequencies of 34.62%, 45.73%, and 35.63% in alpha, beta, and omicron, respectively. In the delta VOC, the mutational landscape differed significantly. The prevalence of T77A, V149A, and T181I was 66.87%, 29.45%, and 10.83%, respectively (Figure S5B). Over the study period, the T77A mutation frequency increased from 12.13% to 80.06%, while V149A and T181I mutation frequencies decreased from 75.53% to 19.25% and 54.54% to 10%, respectively (Figure S5D). T77A was present in all top four sub-lineages of the delta variant, while V149A and T181I were exclusive to B.1.617.2 (Figure S5C).

The S106 - F108 deletion mutation was specific to the BA.2 and BA.2.10 sub-lineages of the omicron VOC, with frequencies of 21.78% and 20.13%, respectively. F108L predominated in all top sub-lineages of the omicron VOC, except for BA.1.1 (Figure S5E). G107L and I189V were specific to the BA.1.1 sub-lineage, with frequencies of 20.17% and 99.8%, respectively (Figure S5E). S106del was replaced by S106T in BA.2.75, with a frequency of 24.55% during Jul’22 - Sep’22. L260F was uniquely associated with XBB.1.16, with a frequency of 98.54% during Oct’22 - Sep’23 (Figure S5E). Importantly, it was observed that S106 - F108del was linked to enhanced ER-zippering activity.

#### Nsp9 protein

Nsp9 plays a crucial role in viral replication and contributes to the assembly of the replication and transcription complex (RTC) [63]. The L42F mutation was found exclusively in the alpha variant with a frequency of 21.47% (Figure S6). Notably, no mutations were detected in the nsp9 of the Indian population concerning the beta, delta, and omicron variants.

#### Nsp12 protein

Nsp12 plays an important role in RNA synthesis process [22–23]. Noteworthy mutational hotspots identified in nsp12 include P227L, P323L, D358N, V359L, and G671S (Figure 4A). Among these, P323L emerges as the most prevalent mutation within nsp12, exhibiting dominance across all VOC with mutation frequencies exceeding 70%. P227L and G671S, with respective frequencies of 19.77% and 96.71%, display specificity for the alpha and delta variants (Figure 4B). The P323L mutation, surpassing 99% frequency in delta and omicron variants, is consistently present in the top four lineages and the top lineage across individual time scales (Figure 4C-4F). Notably, all major omicron sub-lineages share the P323L mutation, while most reported BA.2.75 and XBB.1.16 variants exhibit G671S mutations akin to the delta variant (Figure 4E-4F). The P323L or P323L/G671S mutation correlates with enhanced stability and enzymatic activity of the RdRp complex, thereby facilitating efficient transmissibility [23].

**Figure 4:**
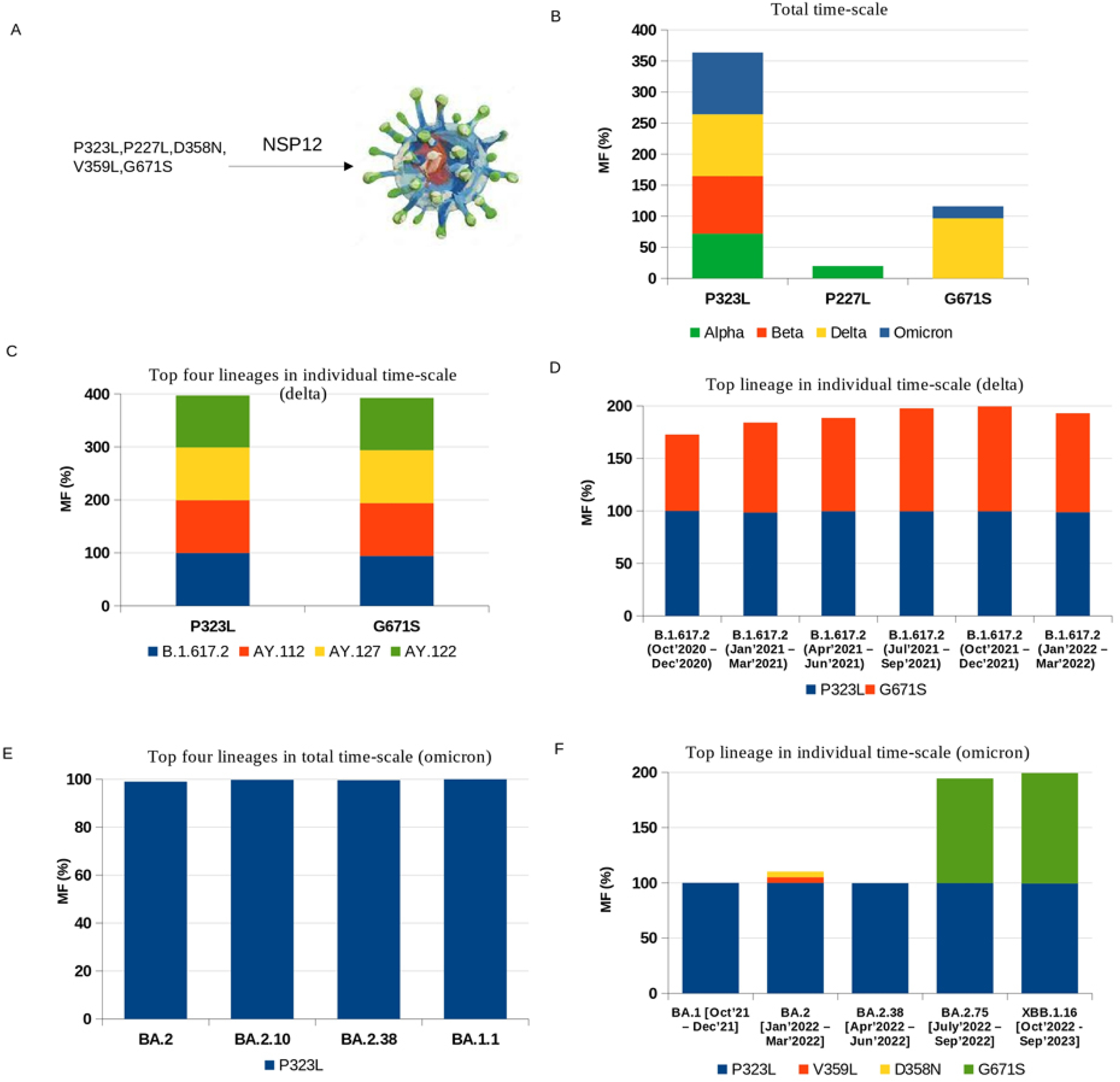
Mutational dynamics of nsp12: A. Mutated amino acids; B. Mutation frequency in total time-scale; C. Mutation frequency of top four lineages of delta variant; D. Mutation frequency of top lineage in individual time-scale of delta variant; E. Mutation frequency of top four lineages of omicron variant; F. Mutation frequency of top lineage in individual time-scale of omicron variant.

#### Nsp13 protein

Nsp13 exhibits both helicase and RNA 5′-triphosphatase activities, engaging in the unwinding of DNA or RNA during RNA replication in an ATP-dependent manner [24]. Notably, prevalent mutations, including S36P, P77A, T588I, I151V, R392C, G170S, and A336T, were observed (Figure S7A). T588I and I151V exclusively occurred in the beta VOC, with mutation frequencies of 17.68% and 9.14%, respectively. The delta variant prominently featured the P77L mutation, among the most frequently observed worldwide (Figure S7B-S7D). In the omicron variant, R392C and S36P were predominant, with frequencies of 81.45% and 25.9%, respectively (Figure S7B). R392C overwhelmingly prevailed, exceeding 94% frequency in all major sub-lineages except BA.1 (Figure S7E). S36P was unique to BA.2.10 and XBB.1.16 (Oct’22 - Sep’23), with a frequency surpassing 99% (Figure S7F).

#### Nsp14 protein

Nsp14 is mainly responsible for proofreading activity [25]. The prevalent mutations of nsp14 were I42V, P46L, V182I, D222Y, and A394V (Figure 5A). It was observed that nsp14 in the Indian population did not exhibit any mutation in the alpha and beta variant. A394V and P46L were specific for delta whereas I42V and D22Y were observed in omicron (Figure 5B). In delta VOC, A394V was prevalent in all top sub-lineages whereas P46L was specific for B.1.617.2 (Figure 5C). In every time-scale, A394V mutation frequency increased from 12.13% to 80.39% (Figure 5D). In omicron VOC, all top four lineages and the top lineage of the individual time-scale had the I42V mutation, which had a >87% frequency (Figure 5E-5F). Interestingly, D222Y was solely specific for XBB.1.16 with a frequency of 99.58% (Figure 5F).

**Figure 5:**
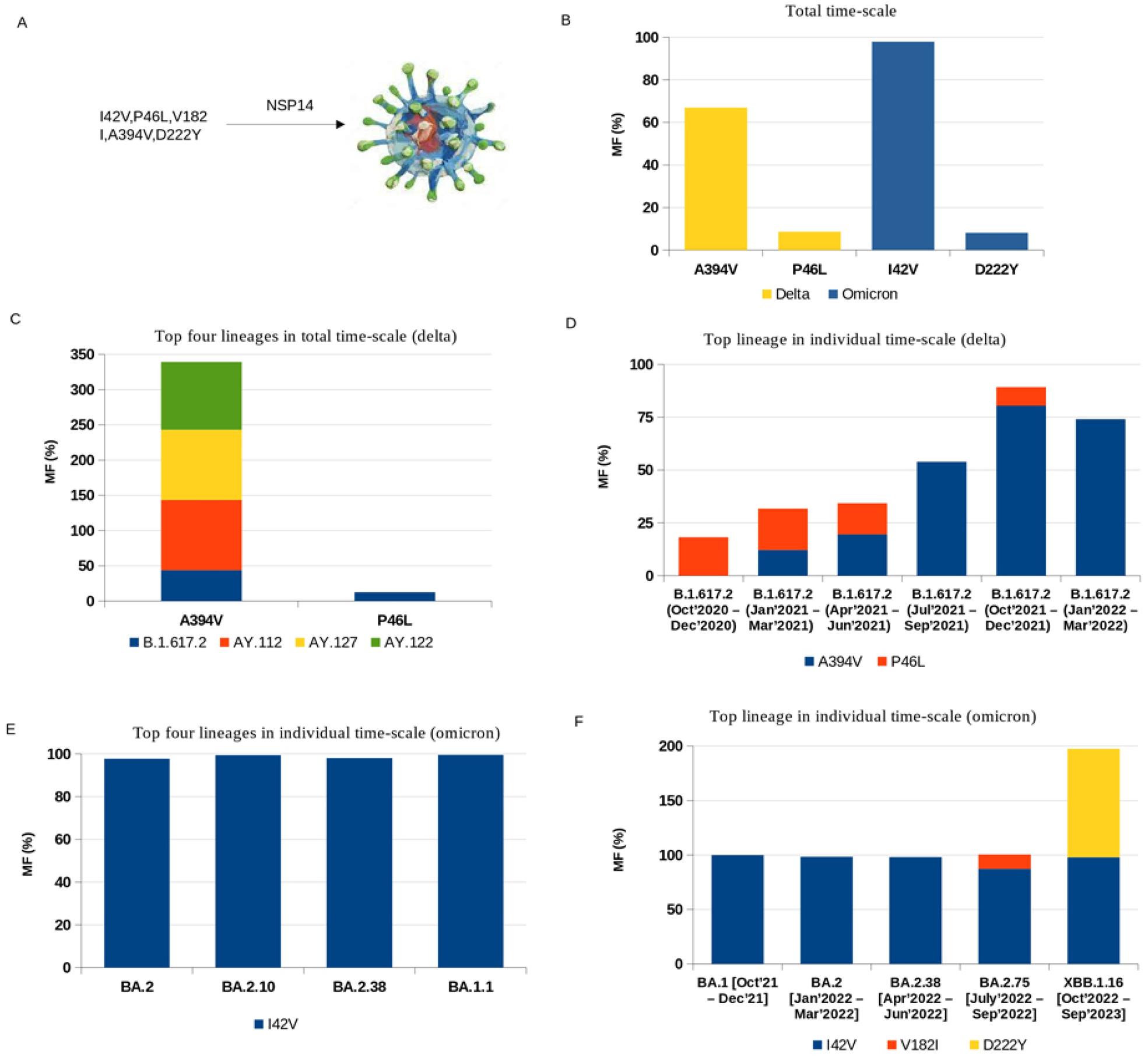
Mutational dynamics of nsp14: A. Mutated amino acids; B. Mutation frequency in total time-scale; C. Mutation frequency of top four lineages of delta variant; D. Mutation frequency of top lineage in individual time-scale of delta variant; E. Mutation frequency of top four lineages of omicron variant; F. Mutation frequency of top lineage in individual time-scale of omicron variant.

#### Nsp15 protein

Nsp15, a conserved uridine-specific endoribonuclease identified in coronaviruses, plays a pivotal role in controlling the viral RNA accumulation and suppressing dsRNA-activated antiviral responses within host cells [64]. Consequently, it stands as a crucial target for the development of both medications and attenuated vaccines. The mutations T112I and H234Y exhibit frequencies of 77.91% and 6.79%, respectively, and are exclusive to the omicron and delta variants (Figure S8A-S8B). Additionally, H234Y and K259R are specific to the B.1.617.2 variant (Figure S8C-S8D). The T112I mutation is prevalent across all major sub-lineages of the omicron VOC, excluding BA.1.1 (Figure S8E-S8F).

#### spike

The mutation frequency of amino acids in the spike protein of SARS-CoV-2 variants of concern in India has exhibited significant variation, especially in regions critical for virus-host interactions, such as the receptor-binding domain (RBD). We calculated and analyzed the mutation frequency of the spike protein over the entire time-scale. Key mutations, including D614G, N501Y, E484K, and L452R, have been frequently observed across VOCs like alpha, beta, delta, and omicron. Specifically, five, six, four, and ten mutations with frequencies greater than 95% were present in the alpha, beta, delta, and omicron variants, respectively (Figure S9). Throughout the study period, D614G was the most prevalent mutation in the alpha and delta variants. However, A701V and H655Y were the predominant mutations in the beta and omicron variants, respectively (Figure S9).

### Phylodynamics of spike and NSPs based on genome and mutational profiling

Next, we wanted to investigate whether mutation profile of individual genes/proteins of the SARS-Cov-2 virus played any significant role in delineating the phylodynamics of various VOC. In order to do that we first created a species wise phylogenetic profile of the various VOC genomes reported and submitted to GISAID database from India. 500 genome sequences were randomly selected representing four VOC (alpha, beta, delta, and omicron, respectively) and ten lineages from the GISAID database where each lineage contains 50 sequences. Genome wide multiple sequence alignment was generated followed by construction of phylogenetic tree using multiple tree generating tools and techniques (Figure 6A). Further, a lineage specific tree was created following the clustering patterns of the individual sub-lineages within each VOC (Figure 6B). Hence, Figure 6B provides a phylogenetic tree for the ten lineages belonging to the four highly prevalent VOC in India.

**Figure 6:**
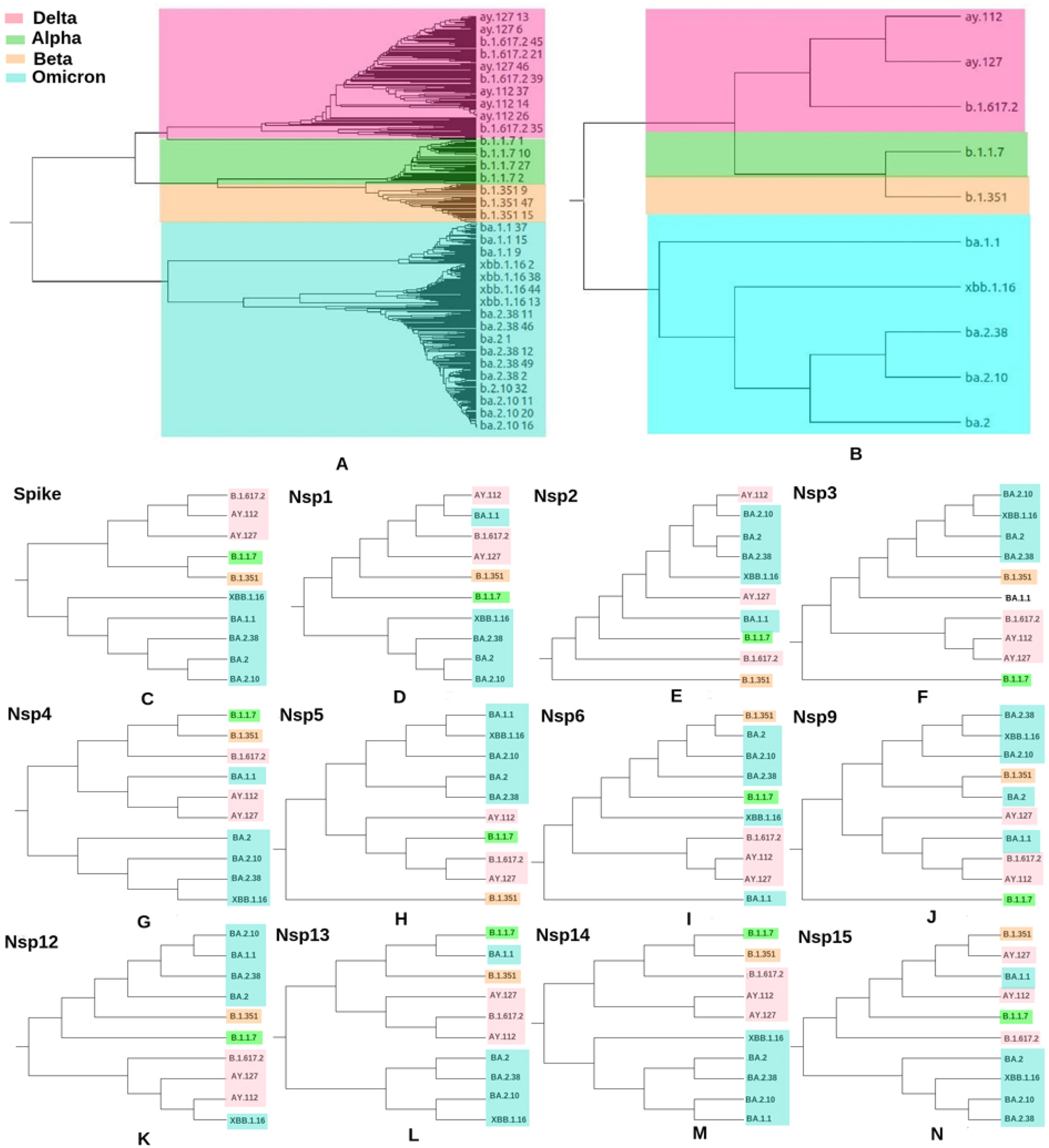
Comparative phylodynamic analysis of VOC using genome-based and mutation-based profiles. A. Phylogenetic tree of VOC. Ten lineages were used to generate the tree. One from alpha (B.1.1.7), one from beta (B.1.351), top tree lineages from delta (B.1.617.2, AY.112, AY.127) and top five lineages from omicron (BA.1, BA.2, BA.2.10, BA.2.38 and XBB.1.16); B. A more simplified linear tree shows a subset of the lineages or variants. Cladograms were generated from the mutational profile of the spike and non-structural proteins (NSPs). C. spike; D. nsp1; E. nsp2; F. nsp3; G. nsp4; H. nsp5; I. nsp6; J. nsp9, K. nsp12; L. nsp13; M. nsp14; N. nsp15

Subsequently, we created mutation frequency matrix for eleven NSPs (nsp1-6, nsp9, and nsp12-15, respectively) and spike protein. For a specific gene/protein, frequencies of all observed mutations were calculated for each of the lineages to create the mutation frequency matrix. Finally, a dendogram tree was prepared for each of the NSP and spike protein (Figure 6C-6N). Next, we compared the genome based phylodynamic tree with individual gene specific mutation frequency derived trees using the Phylo.io and Visual Tree cmp tools [38–39]. Similarity of clustering between the two types of tree is reflected by a lower value of Robinson Foulds (RF) distance (Table 1).

**Table 1:** Comparative analysis of phylogenetic trees between genome-based and mutational frequency based trees.

| Proteins | Phylo.io (RF) | VisualTreecmp (RF) |
| --- | --- | --- |
| spike | 3 | 1.5 |
| Nsp1 | 7 | 3.5 |
| Nsp2 | 11 | 5.5 |
| Nsp3 | 7 | 3.5 |
| Nsp4 | 7 | 3.5 |
| Nsp5 | 9 | 4.5 |
| Nsp6 | 9 | 4.5 |
| Nsp9 | 11 | 5.5 |
| Nsp12 | 11 | 5.5 |
| Nsp13 | 7 | 3.5 |
| Nsp14 | 7 | 3.5 |
| Nsp15 | 9 | 4.5 |

Mutation pattern in spike protein was a critical factor in influencing the evolution of different variants of the SARS-Cov-2 virus enabling it to evade host defence mechanism and establishment and persistence of the infectivity. It was encouraging to observe the most similarity between the spike mutation tree and the overall SARS-Cov-2 genome-based phylogentic trees (Table 1). However, comparison between spike mutation based tree and same derived from other NSPs reveals lesser similarity except nsp14 (Table 2). This is indicative of the possibility that nsp14 mutations could also be influencing factors in guiding the genomic diversity of the virus. Mutation pattern from some of the non-structural proteins like nsp1, nsp3, nsp4, nsp13, and nsp15 could also play crucial role in guiding the phylodynamics and emergence patterns of the SARS-Cov-2 variants of concern.

**Table 2:** Comparative analysis of phylogenetic tree between spike and NSPs.

| Proteins | Phylo.io (RF) | VisualTreecmp (RF) |
| --- | --- | --- |
| Nsp1 | 10 | 6 |
| Nsp2 | 14 | 8 |
| Nsp3 | 12 | 7 |
| Nsp4 | 12 | 7 |
| Nsp5 | 12 | 7 |
| Nsp6 | 12 | 7 |
| Nsp9 | 12 | 7 |
| Nsp12 | 12 | 7 |
| Nsp13 | 10 | 6 |
| Nsp14 | 8 | 4 |
| Nsp15 | 14 | 8 |

### Co-mutation propensity of the most frequent mutations

Co-occurrence of mutations within a specific viral genome from VOC is probably connected to its infectivity and sustenance [65]. Hence, type and frequencies of most frequent co-mutations observed in NSPs and spike within each VOC (alpha, beta, delta, and omicron, respectively) were calculated and compared to identify changes in co-mutation patterns/motifs through emergence of various VOC within Indian populations. Figure 7 shows the highest co-mutations patterns consisting of 3-10 mutations within each VOC.

**Figure 7:**
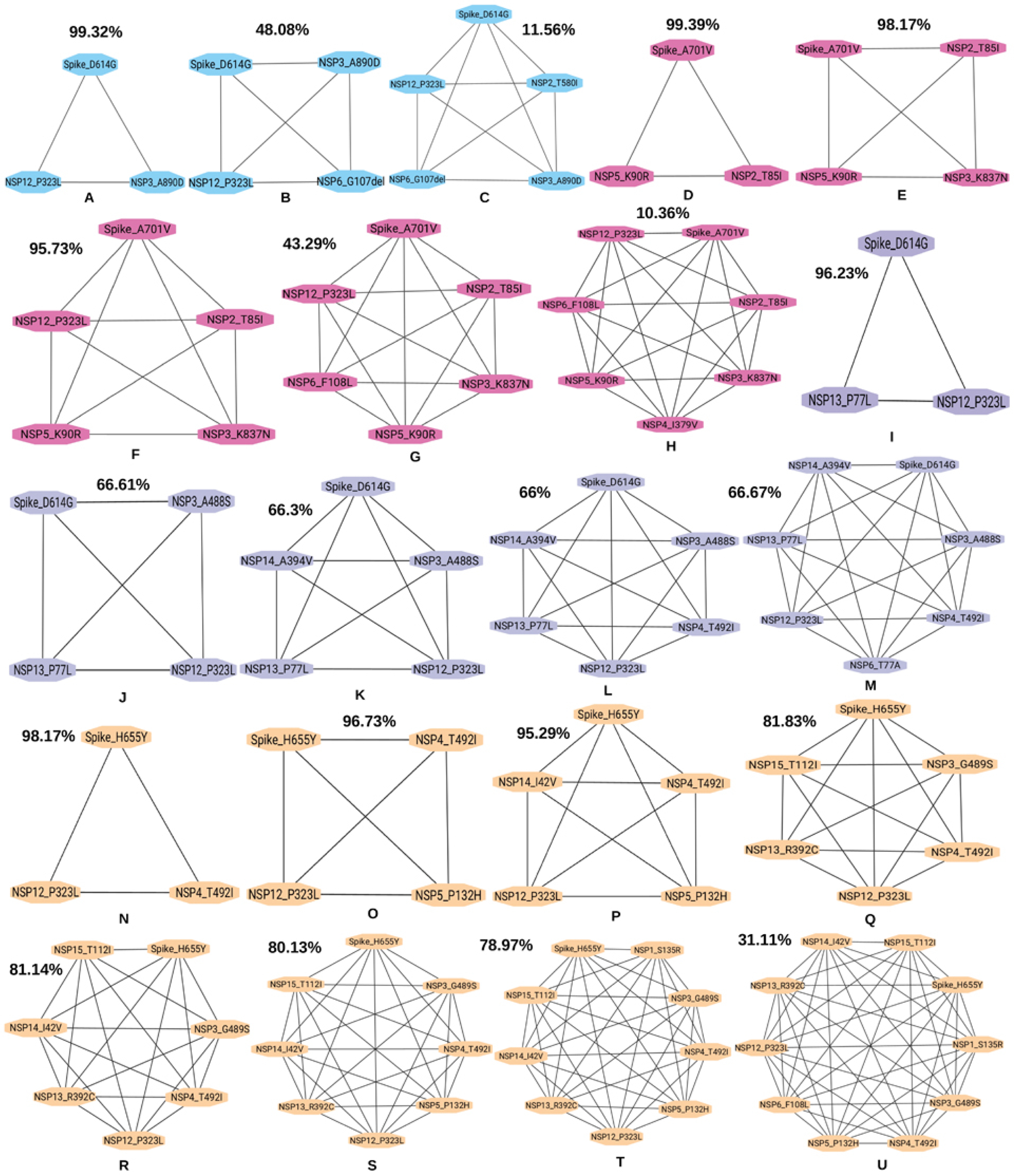
Co-mutation patterns and network of VOC. A-C. 3-5 mutations combination of alpha variant; D-H.3-7 mutations combination of beta variant; I-M.3-7 mutations combination of delta variant; N-U.3-10 mutations combination of omicron variant. Numbers on top of each clique depicts their abundance frequency percentage.

### Structural and energetic alteration of co-mutants across VOC

It is important to investigate whether co-occurrence of the most frequent mutations is responsible for structural and/or functional alterations in the genes/proteins. The cumulative alterations might be responsible for emergence of newer variants with increased infectivity and persistence. Hence, we explored, estimated, and compared the cumulative structural alterations of the co-occurrence combinations of the most frequent mutations across different VOC of the SARS-CoV-2 virus prevalent in India during Oct 2020 to Sep 2023. Three-dimensional (3D) models of the most frequent mutant proteins were generated followed by molecular dynamics (MD) simulations to estimate their structural and energetic alterations resulted due to the mutation. Here, we represented the MD simulation of the five-membered clique (five co-mutations across NSPs and spike) from individual VOC (Table S1-S4). Thousands of intermediate structures (160000 for all 16 mutants) were generated using standard MD algorithms like GROMACS [48] with 100 nano second (ns) simulation for each mutant resulting in altogether 1.6 micro second (μs) MD simulation run. Various structural analyses were performed to study the structural impact of these variants (Table S1-S4). Notably, the average energy value of the five-membered clique of the omicron variant was observed to lower indicating better stability compared to the other variants (Figure 8A and Table 3). Root mean square deviation (RMSD) of the representative mutant structure was calculated with respect to the wild type (WT) structure. For each mutant protein, a representative ensemble structure was selected from the largest cluster generated based on their structural difference reflected by RMSD. The average RMSD was then determined for each co-mutating clique comprising five mutant proteins. The mean RMSD values for the five-membered clique associated with the alpha, beta, delta, and omicron variants were 2.34, 2.32, 2.64, and 2.99 nm, respectively (Figure 8B and Table 3). Similarly, the average fluctuation, as indicated by root mean square fluctuation (RMSF), was evaluated for the most prevalent co-mutational cliques. The average fluctuations observed in the omicron variant were higher than those in the other variants (Figure 8C and Table 3). The radius of gyration (Rg) is a measure of the compactness or spatial distribution of mass within a protein molecule. Mean Rg values of alpha, beta, delta, and omicron were 3, 3, 2.96 and 3.24 nm, respectively (Figure 8D and Table 3). A greater Rg of omicron variant suggests that its protein molecules may adopt more extended conformation due to the presence of more flexible regions or domains that allow it to sample a larger conformational space. The surface areas of the delta and omicron variants were higher, averaging 330.02 nm^2^ and 324.44 nm^2^, respectively, in contrast to 310.44 nm^2^ and 288.90 nm^2^ for the alpha and beta variants (Figure 8E and Table 3).

**Figure 8:**
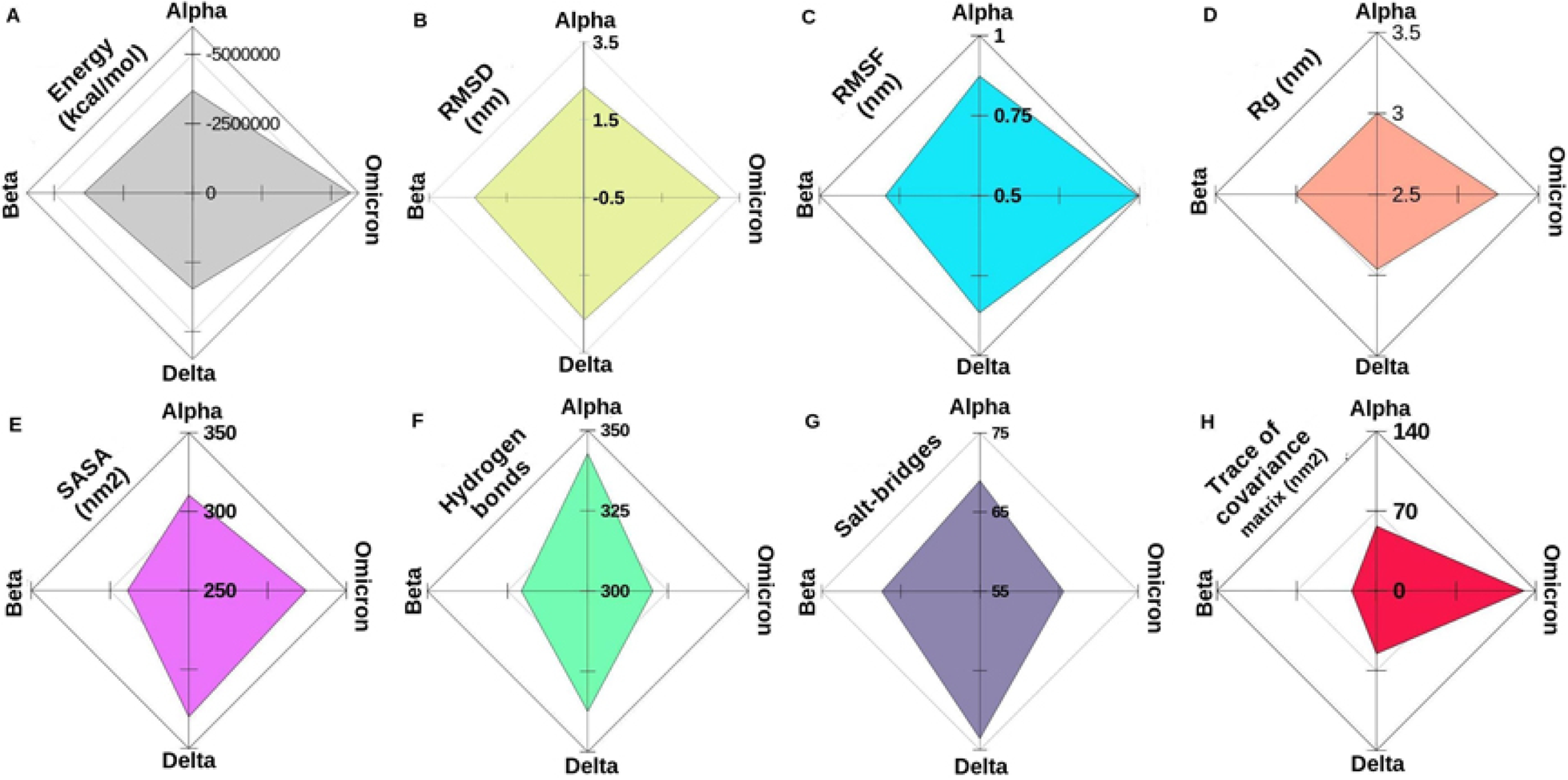
Average MD simulation trajectory parameters of five-membered clique of each VOC. Average values from four VOCs are plotted in four axes. A. Energy; B. RMSD; C. RMSF; D. Rg; E. SASA; F. Hydrogen bonds; G. Salt-bridges; and H. Trace of covariance matrix. [RMSD: Root mean square deviation; RMSF: Root mean square fluctuation; Rg: Radius of gyration; SASA: Solvent-accessible surface are]

**Table 3:**
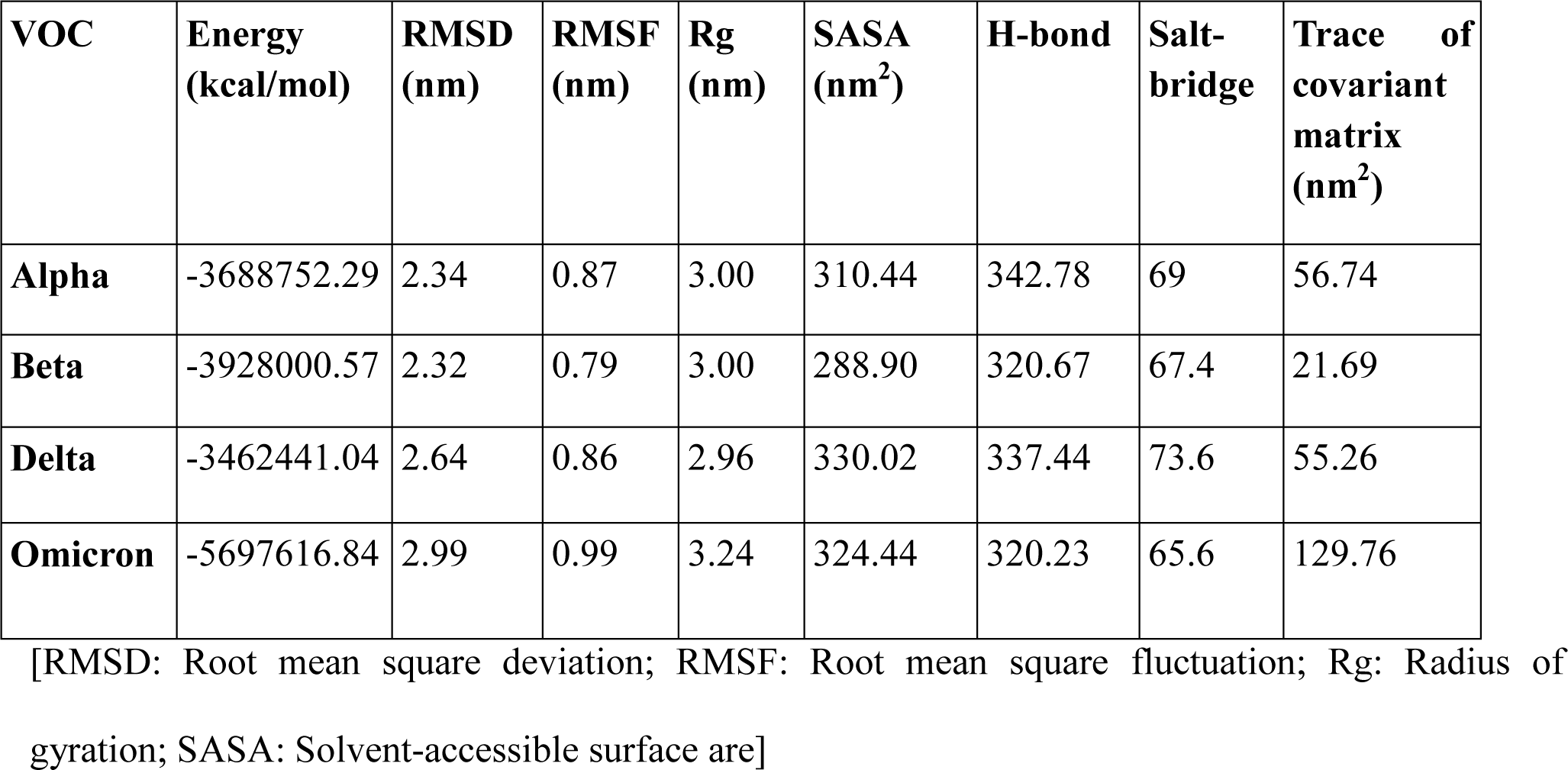
Average trajectory parameter from five-membered clique of each VOC.

| **VOC** | **Energy**<br>**(kcal/mol)** | **RMSD**<br>**(nm)** | **RMSF**<br>**(nm)** | **Rg**<br>**(nm)** | **SASA**<br>**(nm<sup>2</sup>)** | **H-bond** | **Salt-**<br>**bridge** | **Trace of**<br>**covariant**<br>**matrix**<br>**(nm<sup>2</sup>)** |
| --- | --- | --- | --- | --- | --- | --- | --- | --- |
| **Alpha** | -3688752.29 | 2.34 | 0.87 | 3.00 | 310.44 | 342.78 | 69 | 56.74 |
| **Beta** | -3928000.57 | 2.32 | 0.79 | 3.00 | 288.90 | 320.67 | 67.4 | 21.69 |
| **Delta** | -3462441.04 | 2.64 | 0.86 | 2.96 | 330.02 | 337.44 | 73.6 | 55.26 |
| **Omicron** | -5697616.84 | 2.99 | 0.99 | 3.24 | 324.44 | 320.23 | 65.6 | 129.76 |

[RMSD: Root mean square deviation; RMSF: Root mean square fluctuation; Rg: Radius of gyration; SASA: Solvent-accessible surface are]

To explore how mutations within the variants cause the observed changes in the network of long-range correlated motions, we examined alterations in the count of hydrogen bonds and salt-bridges. These changes signify the specificity of intermolecular interactions between secondary structures. The average numbers of intramolecular hydrogen bonds for alpha, beta, delta, and omicron were 342.78, 320.67, 337.44, and 320.23, respectively (Figure 8F and Table 3). Similarly, the mean counts of salt bridges for alpha, beta, delta, and omicron were 69, 67.4, 73.6, and 65.6, respectively (Figure 8G and Table 3). These analyses of interactions suggest that the omicron variant displayed lower average hydrogen bonds and salt bridges, indicating greater flexibility and increased freedom of movement compared to the other variants.

The process of applying principal components analysis (PCA) to a protein trajectory is called Essential Dynamics (ED), which extracts the essential motions from the molecular dynamics (MD) trajectory of specific protein molecules. This analysis was utilized to examine the conformational space and transitions within variants of concern (VOC). The trace values, representing the sum of the eigenvalues, were 56.74 nm^2^, 21.69 nm^2^, 55.26 nm^2^, and 129.76 nm^2^ for the alpha, beta, delta, and omicron variants, respectively. A greater trace value indicated that the proteins of the omicron variant appeared to explore a larger conformational space due to increased flexibility compared to other variants. This observation correlates with the trajectory of other parameters such as RMSD, RMSF, Rg, and intramolecular interactions. Table 4 shows the percentages of average secondary structure of each VOC. It was observed that the alpha-helix content was higher in alpha and beta variants whereas the percentage of beta-sheets or beta-ladder was higher in omicron variant. The loop or unstructured region was higher in delta and omicron variant compared with other variants. Furthermore, random structures such as pi-helix, kappa-helix, 3_10_-helix, turn, and bend also varied in different VOC.

**Table 4:** Average secondary structure parameters from five-membered clique of each VOC.

| **Average**<br>**secondary**<br>**structural**<br>**content (%)** | **Alpha** | **Beta** | **Delta** | **Omicron** |
| --- | --- | --- | --- | --- |
| **Alpha-helix** | 28.06 | 27.28 | 20.67 | 21.14 |
| **Beta-bridge** | 1.14 | 1.36 | 1.73 | 1.8 |
| **Beta-ladder** | 23.78 | 23.83 | 23.33 | 24.52 |
| **3_10-helix** | 3.01 | 3.23 | 3.99 | 3.97 |
| **pi-helix** | 0.69 | 0.48 | 0.38 | 0.69 |
| **Kappa-helix** | 1.1 | 1.34 | 2.11 | 2.15 |
| **bend** | 10.96 | 10.59 | 12.4 | 10.94 |
| **Turn** | 13.06 | 13.18 | 12.77 | 14.18 |
| **Loop** | 18.18 | 18.71 | 22.6 | 21.69 |

## Conclusion

This study aims to elucidate the evolution of SARS-CoV-2 in India by analyzing 432 lineages representing the country’s diverse clades. Utilizing available datasets of VOC, we estimated the frequency of lineages to understand the evolution of the primary lineages and sub-lineages of SARS-CoV-2. A time series analysis was performed to examine the frequency of significant mutations in the circulating SARS-CoV-2 genomes. We conducted an in-depth examination of the distribution of mutations across defined lineages and compared frequently co-occurring mutation pairs. Our genome-based and mutation-based phylodynamics study suggests a strong association between nsp14 and the spike protein. We employed modeling and molecular dynamics simulations of the viral spike and NSP proteins of the VOC to investigate the impact of mutations on protein stability and flexibility. MD simulations and structural analysis of the VOC revealed clear differences between the omicron variant and the alpha, beta, and delta variants. These findings have significant implications for understanding the evolutionary trends of SARS-CoV-2. This research not only enhances our understanding of SARS-CoV-2 but also provides a framework for studying other viral pathogens. The insights gained could drive the development of antivirals and improve our preparedness for future pandemics.

## Acknowledgements

SC acknowledges CSIR-IICB for infrastructure support. AC acknowledges ICMR for post-doctoral Research Associateship [BMI/11(55)2022]. SD thanks DBT for research grant (BT/PR40137/BTIS/137/35/2022) and post-doctoral fellowship.

## Declaration of competing interest

The authors declare that they have no known competing financial interests or personal relationships that could have appeared to influence the work reported in this paper.

## Notes

### Competing Interest Statement

The authors have declared no competing interest.

### Summary of Updates

Some textual correction has been made.

