## Supplementary Materials for "Mutational and evolutionary dynamics of non-structural and spike proteins from variants of concern (VOC) of SARS-CoV-2 in India"

#### Supplementary Figures

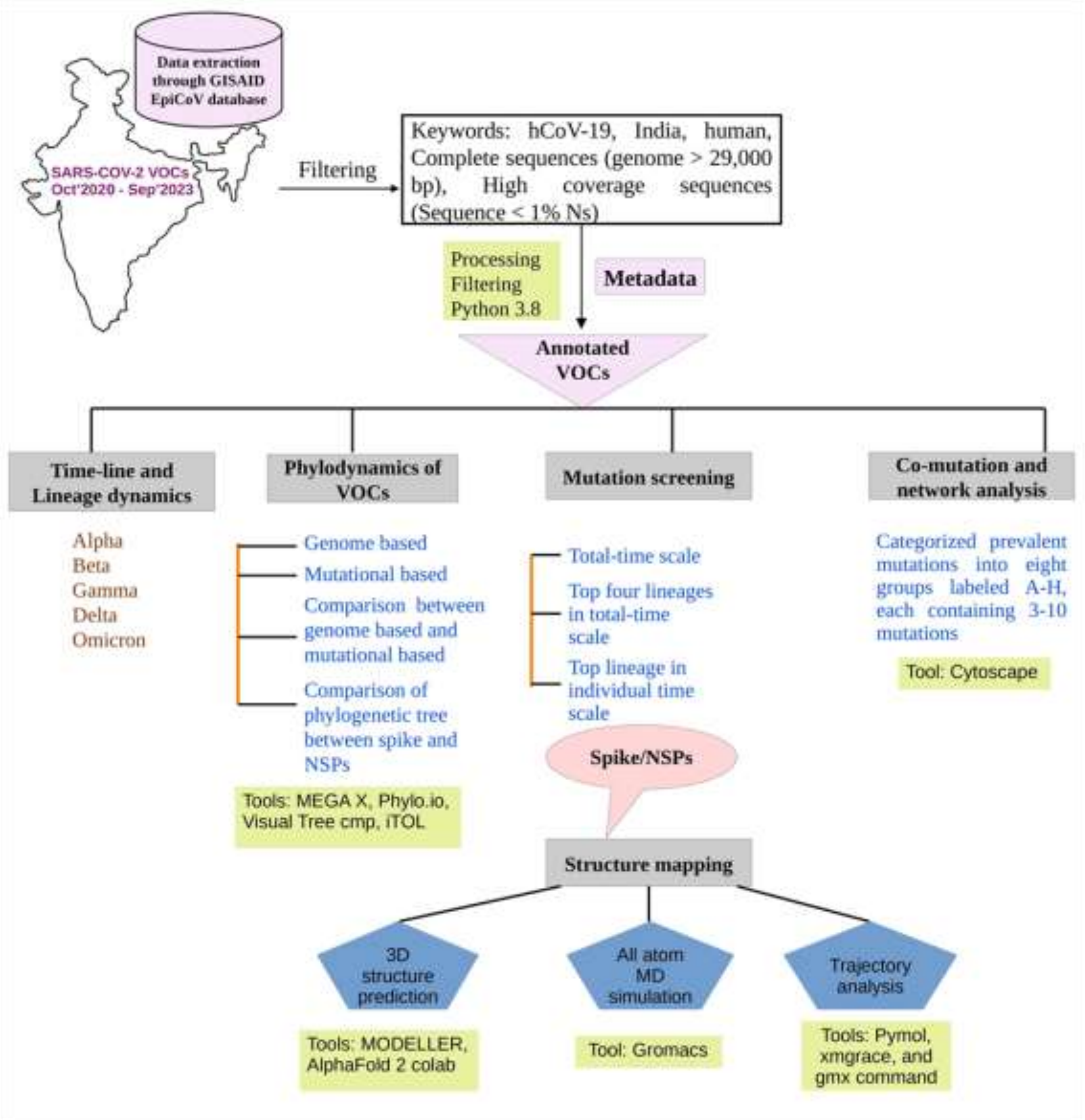


**Figure S1: Flow chart of the overall methodology.**

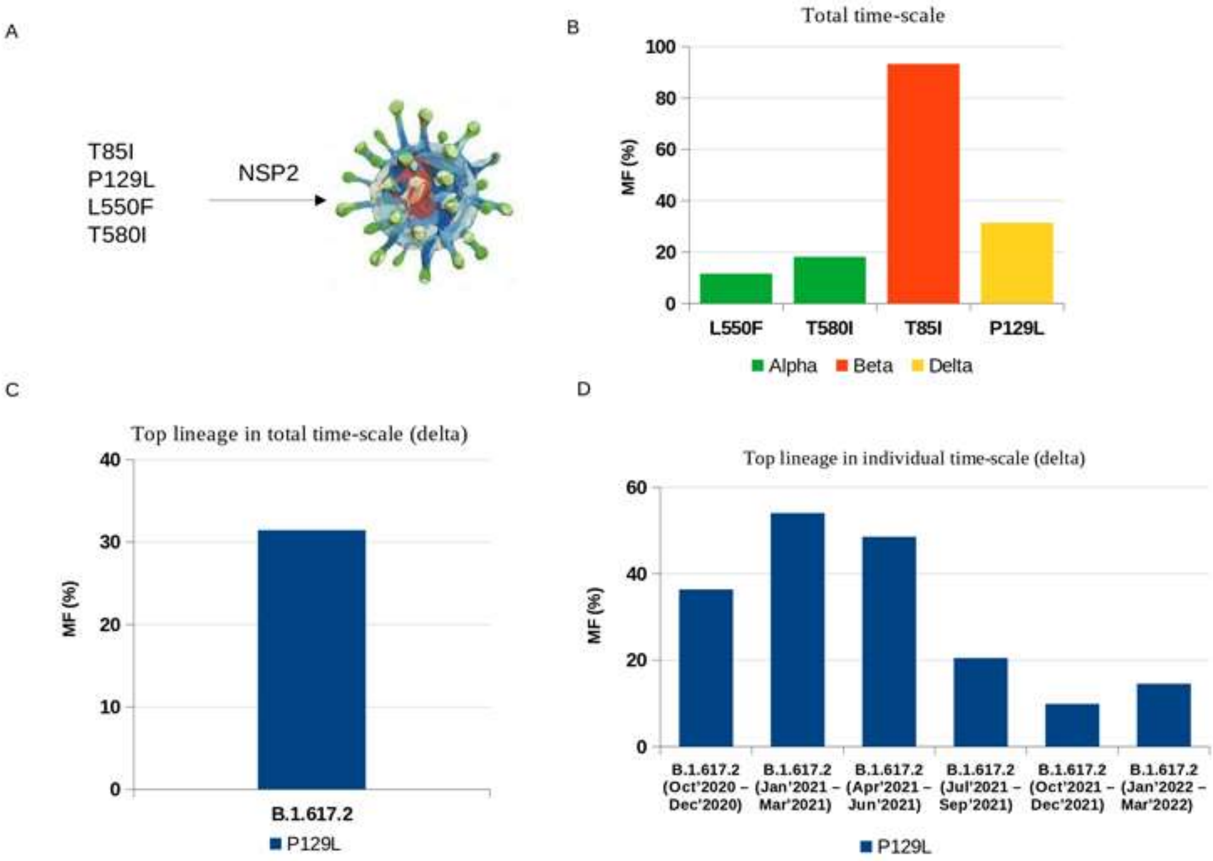


**Figure S2: Mutational dynamics of nsp2:** A. Mutated amino acids; B. Mutation frequency in total time-scale; C. Mutation frequency of top four lineages of delta variant; D. Mutation frequency of top lineage in individual time-scale of delta variant.

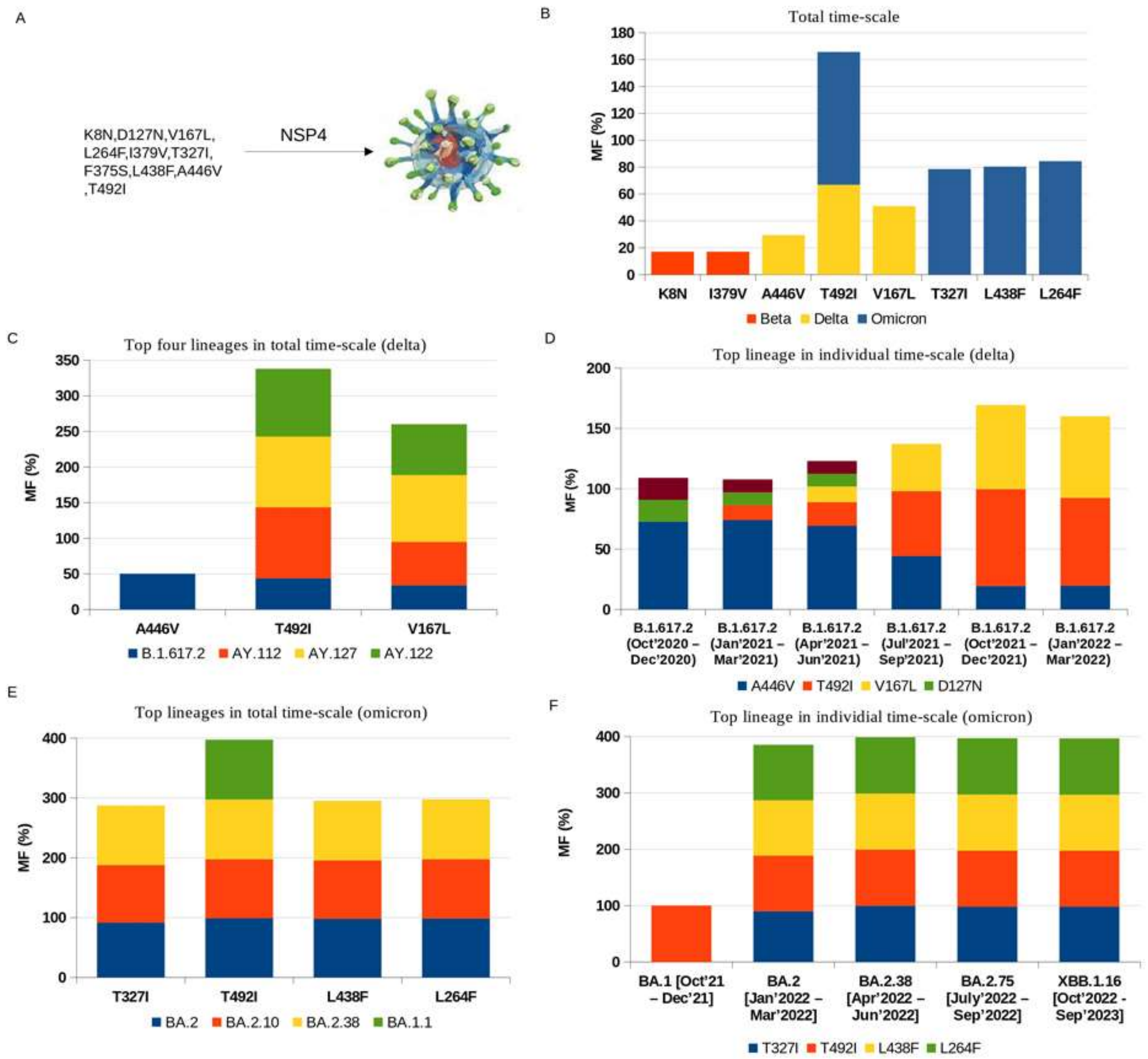


**Figure S3: Mutational dynamics of nsp4:** A. Mutated amino acids; B. Mutation frequency in total time-scale; C. Mutation frequency of top four lineages of delta variant; D. Mutation frequency of top lineage in individual time-scale of delta variant; E. Mutation frequency of top four lineages of omicron variant; F. Mutation frequency of top lineage in individual time-scale of omicron variant

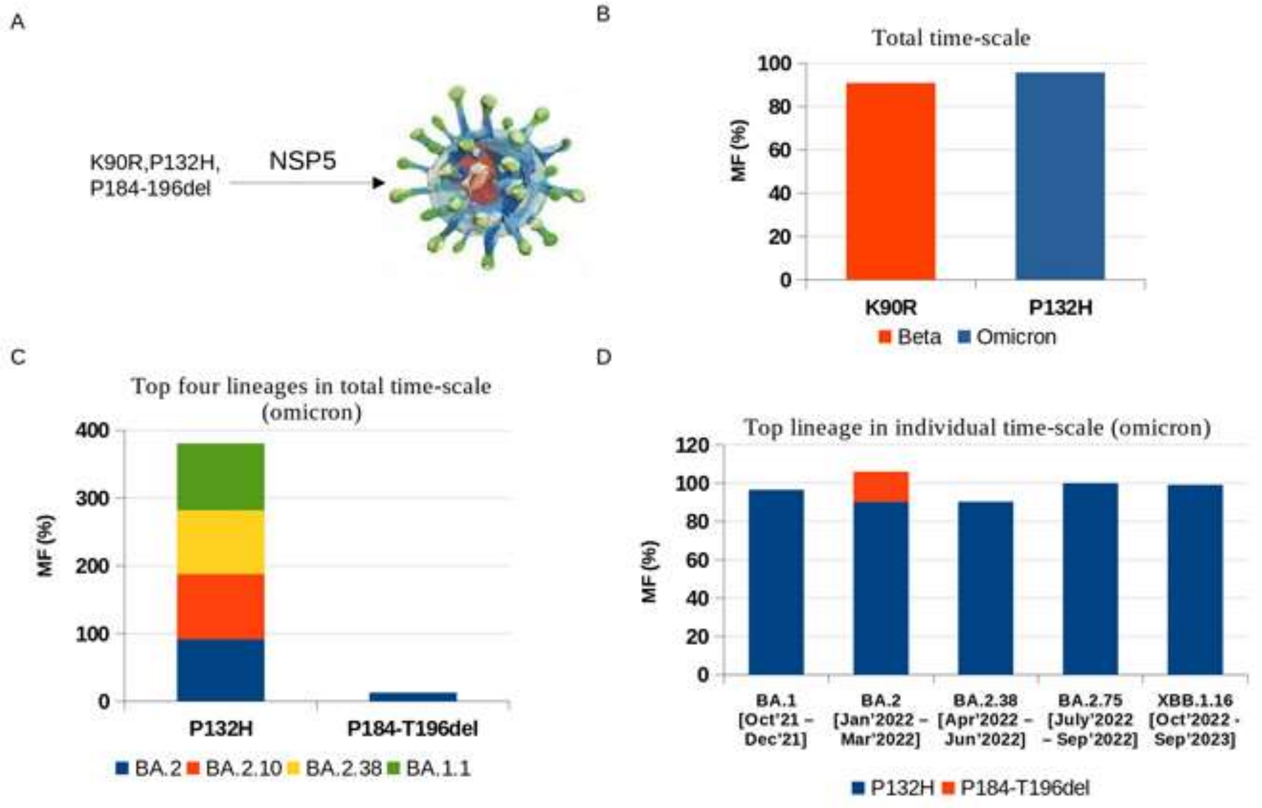


**Figure S4: Mutational dynamics of nsp5**: A. Mutated amino acids; B. Mutation frequency in total time-scale; C. Mutation frequency of top four lineages of omicron variant; F. Mutation frequency of top lineage in individual time-scale of omicron variant

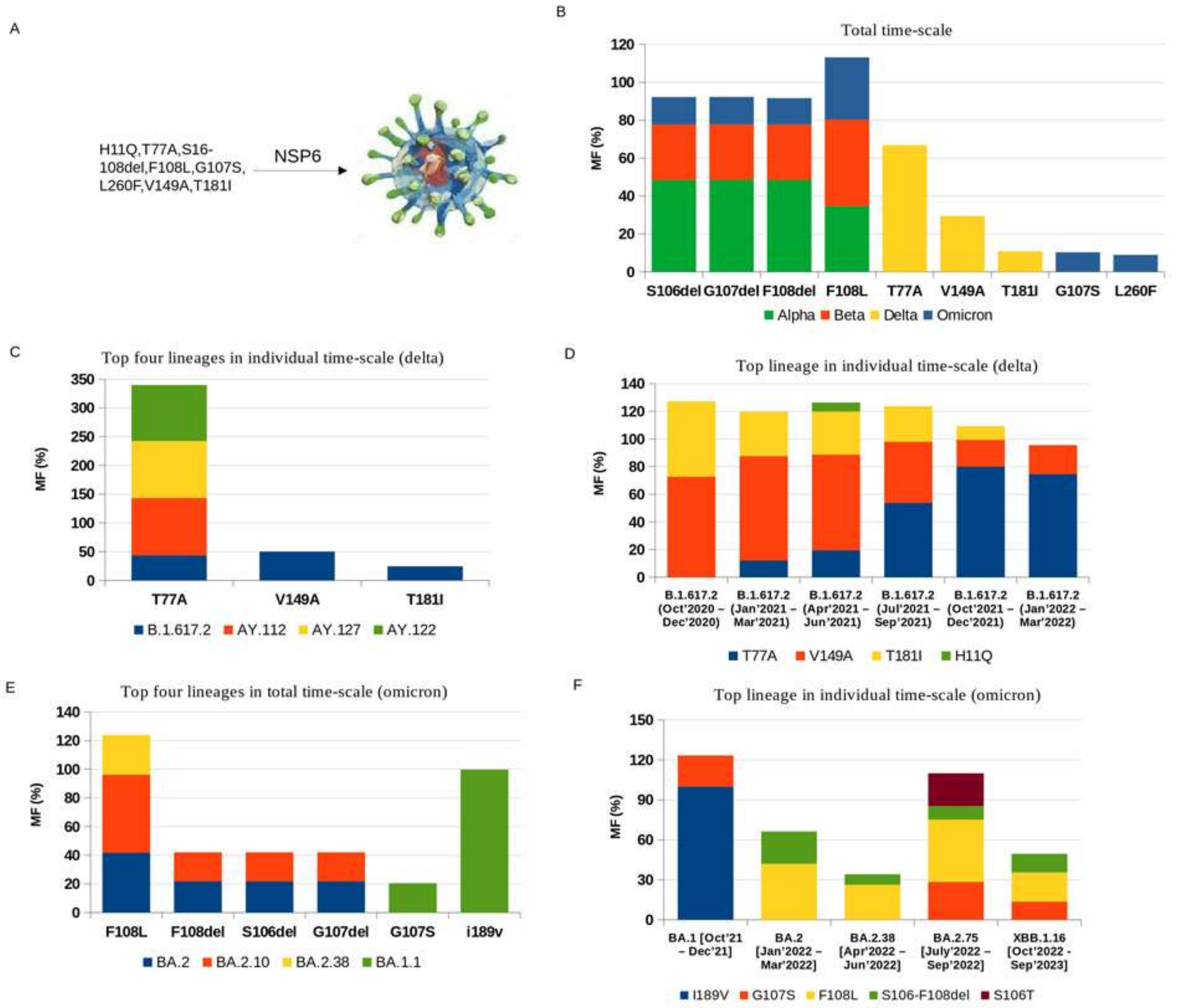


**Figure S5: Mutational dynamics of nsp6:** A. Mutated amino acids; B. Mutation frequency in total time-scale; C. Mutation frequency of top four lineages of delta variant; D. Mutation frequency of top lineage in individual time-scale of delta variant; E. Mutation frequency of top four lineages of omicron variant; F. Mutation frequency of top lineage in individual time-scale of omicron variant.

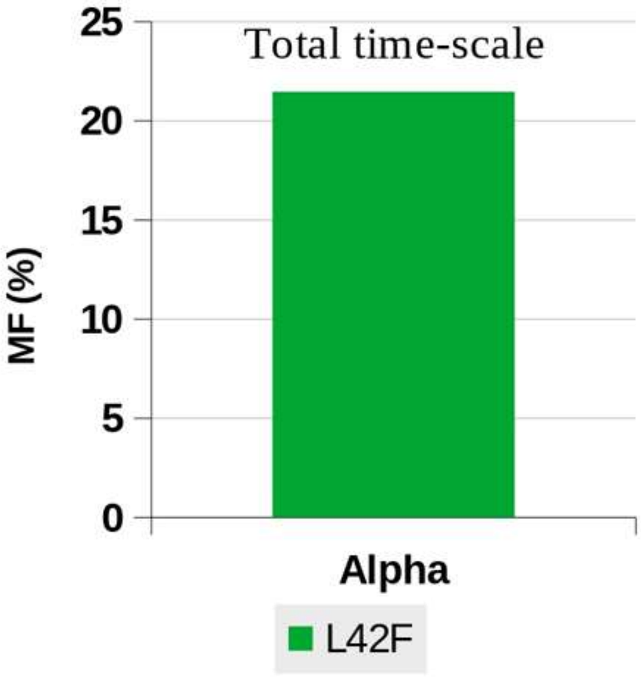


**Figure S6: Mutational dynamics of nsp9:** Most frequent mutation.

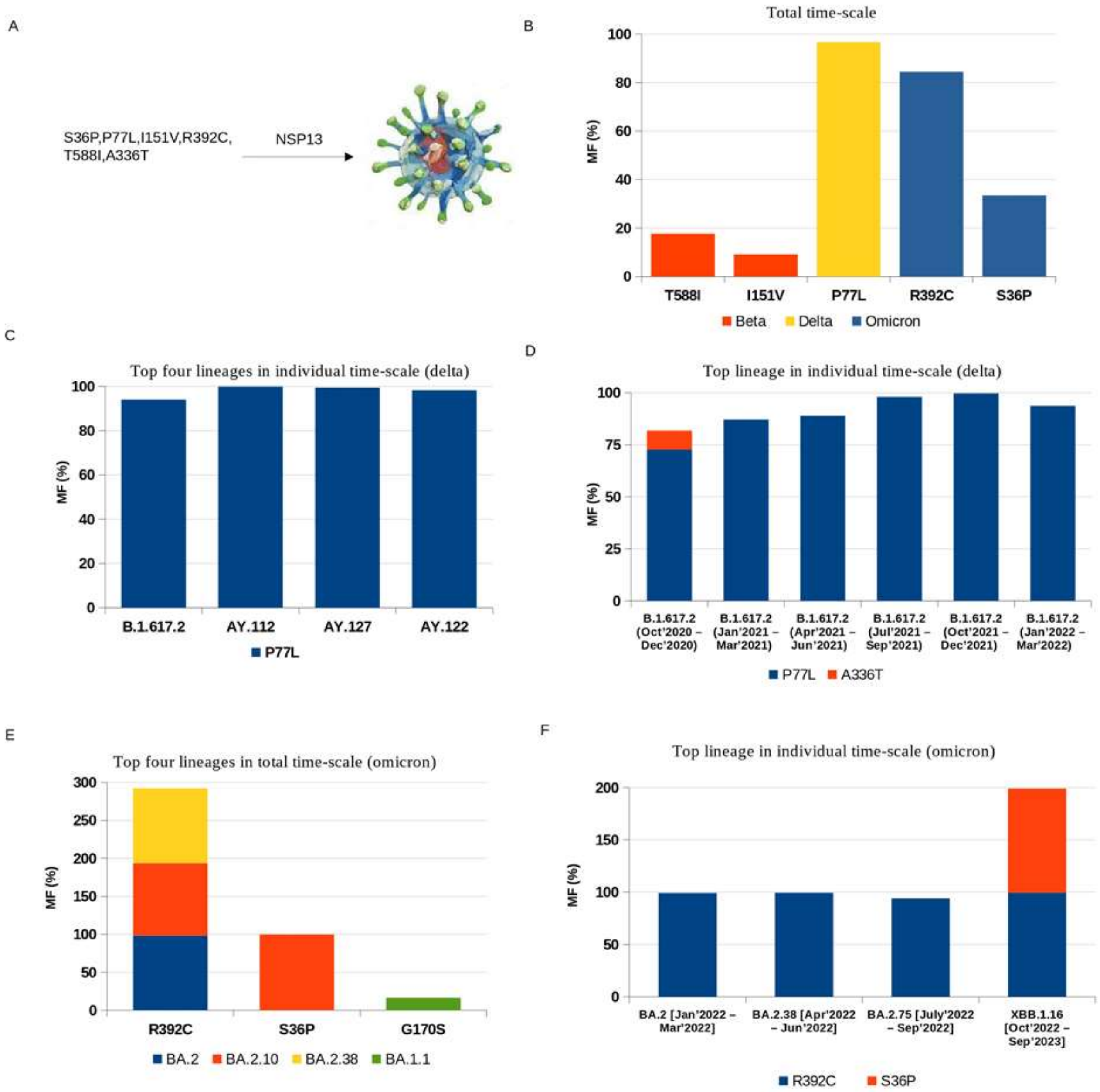


**Figure S7: Mutational dynamics of nsp13:** A. Mutated amino acids; B. Mutation frequency in total time-scale; C. Mutation frequency of top four lineages of delta variant; D. Mutation frequency of top lineage in individual time-scale of delta variant; E. Mutation frequency of top four lineages of omicron variant; F. Mutation frequency of top lineage in individual time-scale of omicron variant.

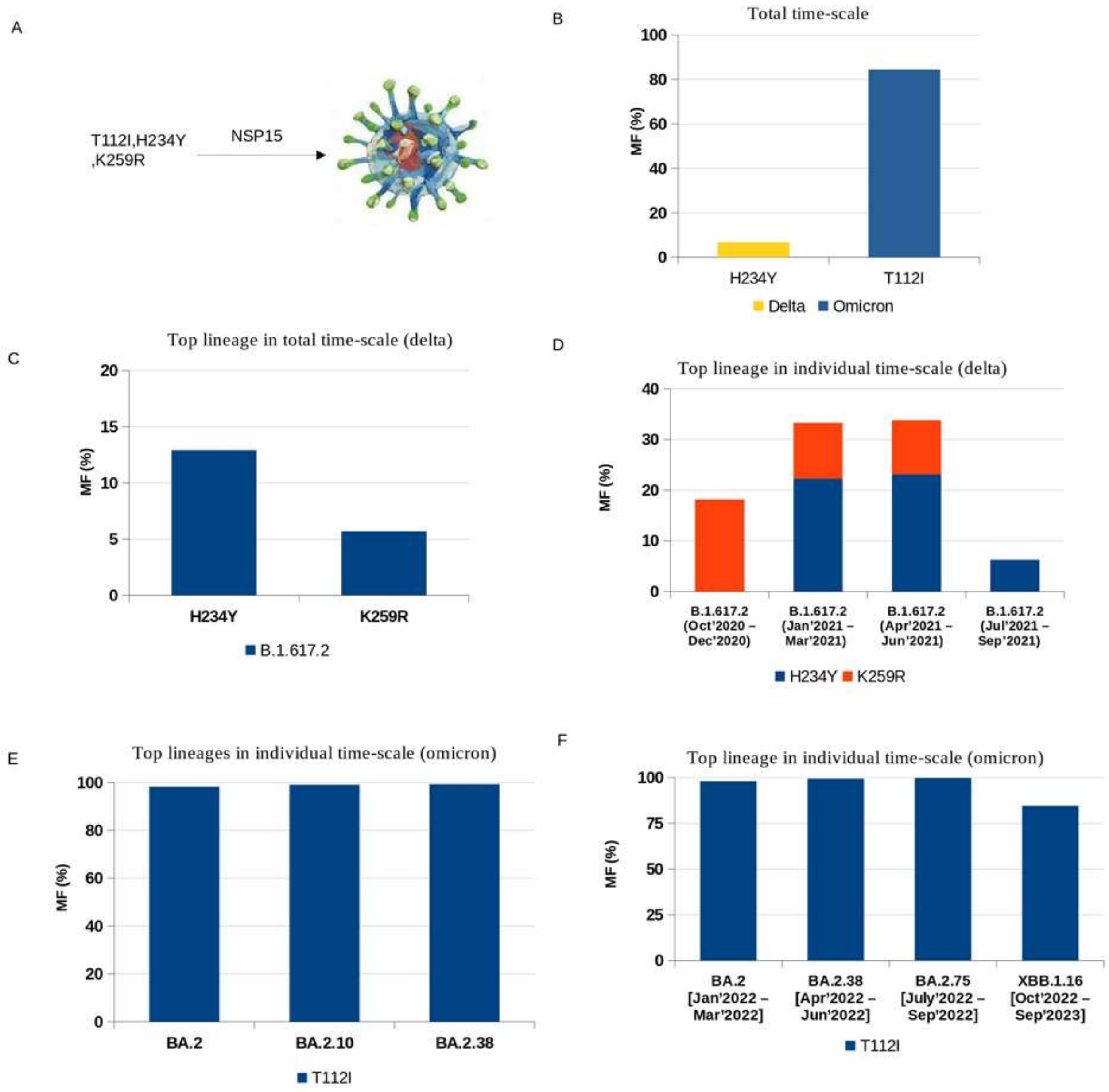


**Figure S8: Mutational dynamics of nsp15:** A. Mutated amino acids; B. Mutation frequency in total time-scale; C. Mutation frequency of top four lineages of delta variant; D. Mutation frequency of top lineage in individual time-scale of delta variant; E. Mutation frequency of top four lineages of omicron variant; F. Mutation frequency of top lineage in individual time-scale of omicron variant.

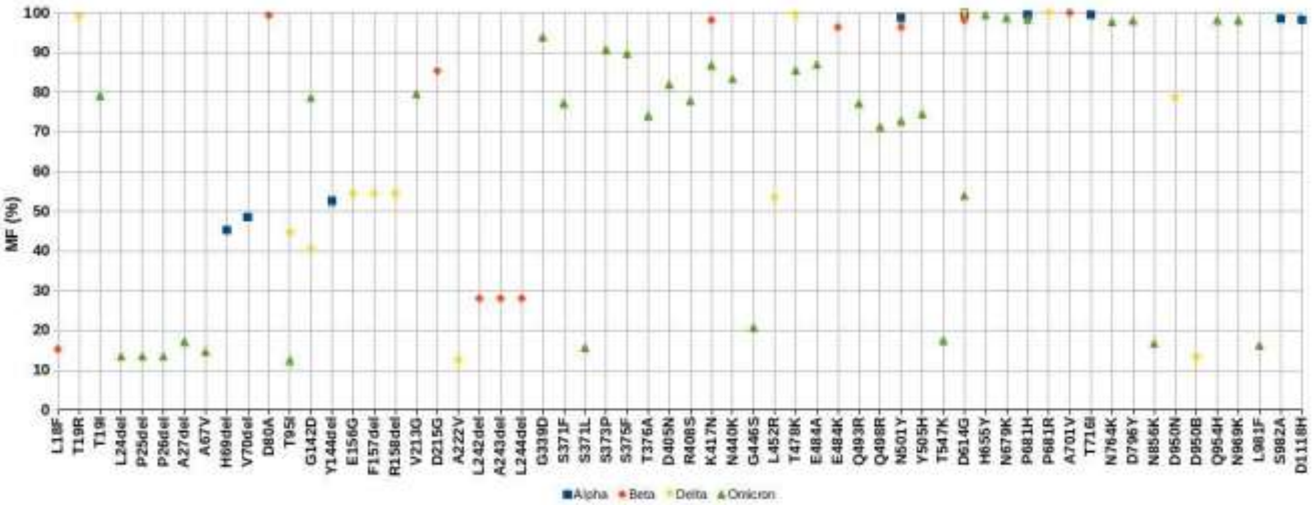


**Figure S9: Mutational dynamics of spike:** Mutation frequency of VOCs in total time scale

### Supplementary Tables

**Table S1: Average trajectory parameters of alpha VOC**

| Five membered clique | RMSD (nm) | RMSF (nm) | Rg (nm) | SASA ($nm^2$) | Energy (kcal/mol) | H-bond | salt-bridge | Trace of covariance matrix |
|---|---|---|---|---|---|---|---|---|
| NSP2_T580I | 0.97 | 0.24 | 3.82 | 345.92 | -6234798.35 | 371.88 | 100 | 42.72 |
| NSP6_G107del | 2.06 | 0.17 | 2.03 | 159.25 | -1050952.59 | 163.06 | 13 | 14.04 |
| NSP12_P323L | 3.34 | 3.34 | 3.25 | 506.07 | -4702064.77 | 589.32 | 142 | 21.34 |
| NSP3_A890D | 1.78 | 0.11 | 2.34 | 165.05 | -1882633.15 | 211.89 | 36 | 5.07 |
| Spike_D614G | 3.56 | 0.51 | 3.56 | 375.95 | -4573312.63 | 377.78 | 54 | 200.56 |
| **Average** | **2.34** | **0.87** | **3** | **310.45** | **-3688752.3** | **342.79** | **69** | **56.74** |

**Table S2: Average trajectory parameters of beta VOC**

| Five membered clique | RMSD (nm) | RMSF (nm) | Rg (nm) | SASA ($nm^2$) | Energy (kcal/mol) | H-bond | salt-bridge | Trace of covariance matrix |
|---|---|---|---|---|---|---|---|---|
| NSP2_T85I | 1 | 0.23 | 3.89 | 345.05 | -6232596.04 | 344.46 | 95 | 37.55 |
| NSP12_P323L | 3.34 | 3.34 | 3.25 | 506.07 | -4702064.77 | 589.32 | 142 | 21.34 |
| Spike_A701V | 3.34 | 0.22 | 3.35 | 257.82 | -5365056.5 | 257.24 | 46 | 42.2 |
| NSP3_K837N | 1.77 | 0.11 | 2.34 | 166.15 | -1888280.02 | 195.99 | 32 | 4.54 |
| NSP5_K90R | 2.17 | 0.08 | 2.2 | 169.45 | -1452005.56 | 216.38 | 22 | 2.86 |
| **Average** | **2.32** | **0.79** | **3** | **288.91** | **-3928000.58** | **320.68** | **67.4** | **21.7** |

**Table S3: Average trajectory parameters of delta VOC**

| Five membered clique | RMSD (nm) | RMSF (nm) | Rg (nm) | SASA ($nm^2$) | Energy (kcal/mol) | H-bond | salt-bridge | Trace of covariance matrix |
|---|---|---|---|---|---|---|---|---|
| NSP3_A488S | 0.84 | 0.12 | 1.97 | 137.9 | -972174.14 | 129.75 | 26 | 3.76 |
| NSP12_P323L | 3.34 | 3.34 | 3.25 | 506.07 | -4702064.77 | 589.32 | 142 | 21.34 |
| NSP13_P77L | 2.57 | 0.17 | 2.84 | 334.45 | -2431712.07 | 329.75 | 80 | 23.04 |
| NSP14_A394V | 2.92 | 0.2 | 3.2 | 295.74 | -4632941.6 | 260.64 | 66 | 27.64 |
| Spike_D614G | 3.56 | 0.51 | 3.56 | 375.95 | -4573312.63 | 377.78 | 54 | 200.56 |
| **Average** | **2.64** | **0.87** | **2.96** | **330.02** | **-3462441.04** | **337.45** | **73.6** | **55.27** |

**Table S4: Average trajectory parameters of omicron VOC**

| Five membered clique | RMSD (nm) | RMSF (nm) | Rg (nm) | SASA ($nm^2$) | Energy (kcal/mol) | H-bond | salt-bridge | Trace of covariance matrix |
|---|---|---|---|---|---|---|---|---|
| NSP4_T492I | 2.99 | 0.86 | 4.02 | 276.62 | -13118974.14 | 121.34 | 45 | 414.34 |

| | | | | | | | | |
|---|---|---|---|---|---|---|---|---|
| NSP12_P323L | 3.34 | 3.34 | 3.25 | 506.07 | -4702064.77 | 589.32 | 142 | 21.34 |
| NSP14_I42V | 2.92 | 0.2 | 3.18 | 294.81 | -4643988.62 | 283.62 | 56 | 27.07 |
| Spike_H655Y | 3.57 | 0.47 | 3.58 | 375.04 | -4572722.1 | 391.13 | 54 | 182.2 |
| NSP5_P132H | 2.15 | 0.1 | 2.23 | 169.66 | -1450334.62 | 215.78 | 31 | 3.88 |
| **Average** | **2.99** | **1** | **3.25** | **324.44** | **-5697616.85** | **320.24** | **65.6** | **129.77** |
